# The energetic cost of human standing balance and gait initiation over a range of natural postures

**DOI:** 10.1101/2025.09.15.676217

**Authors:** Matto Leeuwis, Nikki van Aerts, Ajay Seth, Patrick A. Forbes

**Affiliations:** Department of Neuroscience, Erasmus MC, University Medical Center Rotterdam, Rotterdam, The Netherlands; Department of Biomechanical Engineering, Delft University of Technology, Delft, The Netherlands

**Keywords:** Postural control, Energy expenditure, Metabolic cost, Gait initiation, Musculoskeletal modeling

## Abstract

Humans typically select movements that minimize energetic cost, a principle most clearly observed during locomotion. Whether such optimization of energy expenditure also governs standing balance remains unclear because its energetic cost has not been systematically quantified across a range of natural postures. Moreover, because standing is often the resting state from which most walking begins, the optimization of posture may also reflect the energetic demands of initiating gait. In this study, we use a combination of indirect calorimetry and musculoskeletal simulations to characterize the energetic cost of standing and gait initiation across natural standing postures and investigate whether humans optimize energy expenditure under these conditions. In Experiment 1 (*N* = 13), we measured metabolic cost at preferred and six different prescribed whole-body orientations. Energy expenditure was lowest at a slight anterior orientation (1.15°) and increased monotonically with whole-body angle, rising twice as fast posteriorly compared to anteriorly. This asymmetry challenges the common modeling simplification that effort is symmetric and linear or quadratic with lean angle. Furthermore, participants preferred body orientations (1.50 ± 0.73°) with similar energy expenditure to the minimum-cost orientation but with significantly more postural variability, suggesting that strict postural regulation was not necessary for energy-optimal control. In Experiment 2 (*N* = 20), participants initiated forward and backward walking from preferred or prescribed lean orientations. Participants did not alter their standing posture before expected gait initiations in the forward or backward direction, consistent with musculoskeletal simulations showing that leaning further in the anticipated direction did not significantly improve gait initiation time or energetic costs. Together, these findings suggest that postural strategies optimize energy efficiency when permitted by the demands of movement readiness. Our study quantifies the energetic cost landscape that governs human postural control, challenges widely used inverted pendulum estimations of this cost, and offers an empirical foundation for developing more accurate simulations of posture and energy expenditure.

**Author summary:** Humans are thought to move in ways that save energy. This idea is well supported for walking, but it is not known whether we do the same during quiet standing. Furthermore, because standing is our idle state from which we initiate movement, we may optimize our posture to ease these transitions. In this study, we investigated whether humans stand in postures that minimize energy expenditure. First, we measured and simulated how the cost of posture changes over a range of natural whole-body orientations and determined that humans tend to choose postures close to the orientation with the lowest cost. Leaning backward incurs an additional energetic cost at twice the rate of forward-leaning postures. Second, we investigated whether expecting to walk in the forward or backward direction affects our preferred posture. Surprisingly, participants did not change their posture in preparation for the known direction of walking. Simulations demonstrated that the energetic benefit of doing so was small. Overall, our findings show that maintaining a slight forward lean results in optimal energy expenditure during standing and gait initiation. However, commonly used assumptions of how energy expenditure varies with lean angle do not match the measured cost distribution and those predicted by musculoskeletal simulation.

## Introduction

Human movement control is shaped by the optimization of competing cost functions that account for the efficiency, accuracy, and energy expenditure required to achieve a desired movement goal. In this framework, a controller monitors cost functions and executes a strategy to minimize them. During locomotion, for example, humans and most other animals tend to choose speeds and modes (e.g., running or walking) that minimize energetic cost (1–10). Because the energetic landscape for gait is well characterized using simulations (4, 11–14) and indirect calorimetry measurements (1, 5, 6), it is possible to determine optimal gait patterns (12–14), simulate new ones (6, 10), and even further optimize them with assistive devices (5, 15). Given the strong evidence of energy optimization in normal locomotion, it is compelling to propose that humans also minimize energetic cost while standing. This idea is reflected in many models of quiet standing, which represent postural control as a single-link inverted pendulum actuated by a single moment around the ankle (16–28). In this simple representation, the moment required to maintain a static orientation scales linearly with the ankle angle. Accordingly, models of postural control are commonly formulated as linear controllers (e.g., proportional-derivative control or linear quadratic regulator) that use the effort (i.e., ankle moment) (21, 24, 28) and/or deviations from a single reference angle or posture (21–27) as cost functions. As a result, most inverted pendulum models of posture implicitly assume that the cost function representing effort in standing can be characterized as a linear or quadratic function of lean angle, with a single optimal posture in the center.

The actual metabolic cost of standing balance, however, may not follow this relationship. Human physiology is more suited for leaning in the anterior direction, with a long forefoot and plantarflexor muscles that enable efficient tonic load bearing (16, 29, 30). Furthermore, numerous joints must be stabilized simultaneously (26), with muscles spanning multiple joints and having different functions and metabolic properties (29–31). As a result, energy expenditure may not change linearly with standing angle and is likely asymmetric in the anterior and posterior directions. Although energy expenditure has been measured extensively in preferred posture with and without vision (32–38), and in challenging postural conditions that increase variability (36–38), there is limited direct experimental evidence characterizing the metabolic cost across a range of natural standing postures. Without this characterization, it is not possible to determine whether humans optimize energy expenditure during quiet standing in the same way they do during gait (1–10). Furthermore, the common modeling simplification that the cost function for effort in posture can be described as symmetric and directly proportional to lean angle (21–27) remains unverified.

An important second consideration for optimizing energy expenditure is that, in daily life, static postures often serve as idle states between brief periods of walking. The most common duration between bouts of walking is ∼10 seconds, and 60% of walking intervals last less than 30 seconds (39). Given the abundance of walking transitions from quiet standing, this raises the possibility that the optimization of postural control should also include the energetic cost or time required to initiate movement from that position. For example, athletes begin their sprint from a forward-leaning crouched position, as this enables maximum acceleration. However, this posture is energetically demanding and unsuitable as a resting state. Conversely, sitting expends less energy than standing (32) but is costly for initiating gait. Optimal control in quiet standing could represent a compromise between the cost of static standing and the posture-dependent cost of gait initiation, acting under the constraint of time. Previous work has shown that temporal and kinematic features of forward gait initiation remain relatively stable across whole-body lean angles (40). However, it remains unknown whether these different postures differ in energetic costs for gait initiation, or how these costs might affect backward gait initiation. We hypothesize that postural control during quiet standing reflects the optimization of total metabolic cost that includes both the energetic demands of static balance and those of initiating movement in the expected direction. We further expect this function to be shaped by physiological asymmetries in the human body, resulting in a cost that may not be symmetric or proportional to lean angle.

In this study, we examined (i) how metabolic cost varies with standing posture, (ii) how posture affects the energetic cost and time required to complete gait initiation, and (iii) whether humans optimize posture to minimize these costs during quiet standing and in preparation for walking. In Experiment 1 (*N* = 13), we measured kinematics, metabolic cost, and muscle activity as participants stood at a preferred posture and six prescribed sagittal-plane postures, and simulated energy expenditure using a musculoskeletal model. In Experiment 2 (*N* = 20), we used our validated musculoskeletal model to estimate the metabolic cost of initiating gait from three prescribed postures or each participant’s preferred posture in both forward and backward directions. Our experimental results and musculoskeletal modeling showed that energy expenditure was lowest at a slight forward lean (2 · 10^-2^ rad; 1.15°) and increased asymmetrically, rising more steeply when leaning backward compared to forward. Participants preferred to adopt body orientations (1.50 ± 0.73°) near the energy minimum, but did so with markedly higher postural variability, indicating that precise stabilization at a single orientation was not essential for maintaining low energetic cost. Musculoskeletal simulations further revealed that whole-body orientation only had a minimal effect on the energetic cost and time spent initiating gait. Consistent with this, participants did not adjust their standing posture to the direction of gait initiation, even when the direction was known. Taken together, our results provide the first direct characterization of the energetic landscape of quiet standing and gait initiation over a range of natural postures and suggest that human postural control approximates optimal energy expenditure by operating near, but not strictly at, a slight anterior optimal whole-body orientation.

## Results

### Energetic cost landscape of human standing balance

In our first experiment, we measured participants’ energy expenditure at various postures using indirect calorimetry to determine how energy expenditure varies over natural postures (see Figure 1A and 1C-E). Participants (*N* = 13) performed two blocks of all trials, each containing a set of preferred posture trials (5 min with eyes open and eyes closed, Figure 1C) where participants self-selected a posture, and a set of target trials (2 min at six targets, random order, Figure 1D) where whole-body orientations (−2, 0, 2, 4, 6, 10 · 10^-2^ rad, equal to −1.15, 0, 1.15, 2.29, 3.44, 5.73°) were prescribed using real-time feedback of the center of pressure position (CoP). Whole-body angle was estimated using an inverted pendulum approximation, where the length of the pendulum was approximated as a percentage of body height and the center of mass (CoM) was assumed to be directly above the CoP. The first 60 seconds of calorimetric data were excluded in all trials to ensure that energy expenditure was measured at a steady state (41) (see S1 Appendix for validation), and the energetic cost was normalized to body weight. For visualization, the basal metabolic cost (i.e., the energy expended by non-motor processes to uphold bodily function) was removed by subtracting the cost of the preferred posture trial with eyes open per participant for both the measured and simulated data (Figure 2A; data without this subtraction are shown in the inset, see Materials and Methods).

**Figure 1:**
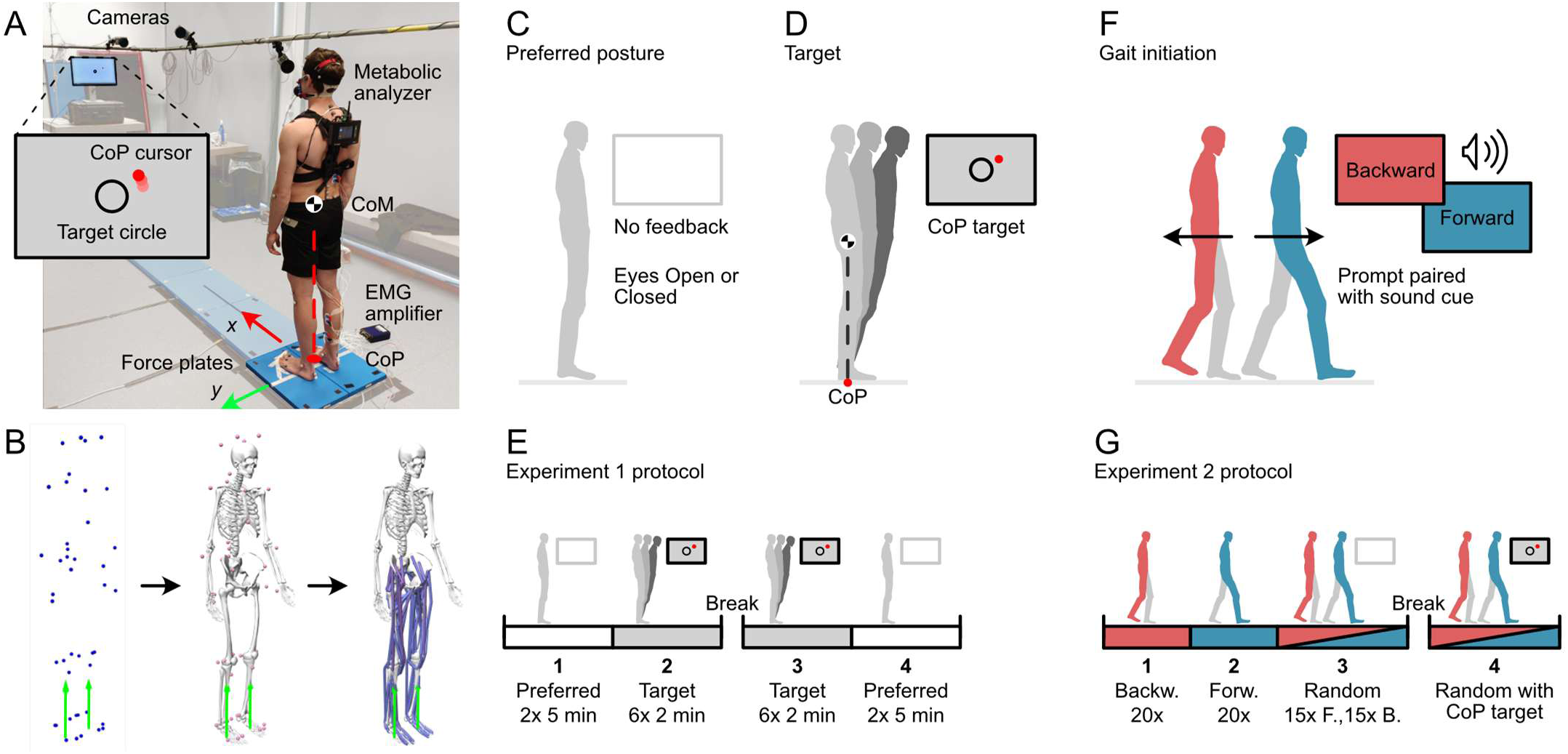
Experimental overview. A: Overview of the setup used in Experiment 1. Participants stood on two force plates and had a total of 43 motion capture markers on their bodies to measure their three-dimensional movement. The back-worn metabolic analyzer measured the flow and air composition at the facemask worn by the participant. The EMG amplifier recorded EMG signals of 8 muscles on the right leg. During target trials, the screen presented a target circle and a cursor of the center of pressure (CoP) in real-time. In Experiment 2, a single force plate was used, and energy expenditure was only estimated from our musculoskeletal model (see Materials and methods). B: Workflow to estimate muscle activation and energy expenditure from motion data. Motion capture data (first panel, blue dots) were used to scale a personalized musculoskeletal model (12) for each participant in OpenSim (13). Inverse kinematics were then performed for all trials to find their joint angles (second panel). The ground reaction forces (green arrows) were then used to calculate the forces and activations in all of the 80 modeled muscles at 20 Hz (third panel). Lastly, the model by Umberger, Gerritsen (30) was used to compute the energy expended by each of the muscles. C: Preferred posture trial; participant self-selected a posture throughout the trial without any feedback on the screen. D: Target trials; whole-body lean angles were prescribed by asking participants to maintain the CoP cursor in a target, which was estimated as the projection of the CoM on the ground plane at the given angle. E: Protocol for Experiment 1. F: Gait initiation trials; participants stood at a preferred posture or at a CoP target, and after 6-10 seconds, the screen depicted a prompt (Forward or Backward) to walk in the corresponding direction. Both prompts were paired with a unique sound cue, and a second tone was played 3.1 s after the first prompt to end the trial. G: Protocol for Experiment 2. In the first block, participants stood at a self-selected preferred posture (panel C) and performed three sets of gait initiation trials (panel E) in random order (Forward, Backward, Random). During the second block, participants were presented with CoP targets (panel D) and received a prompt to walk in the forward or backward direction chosen at random.

**Figure 2:**
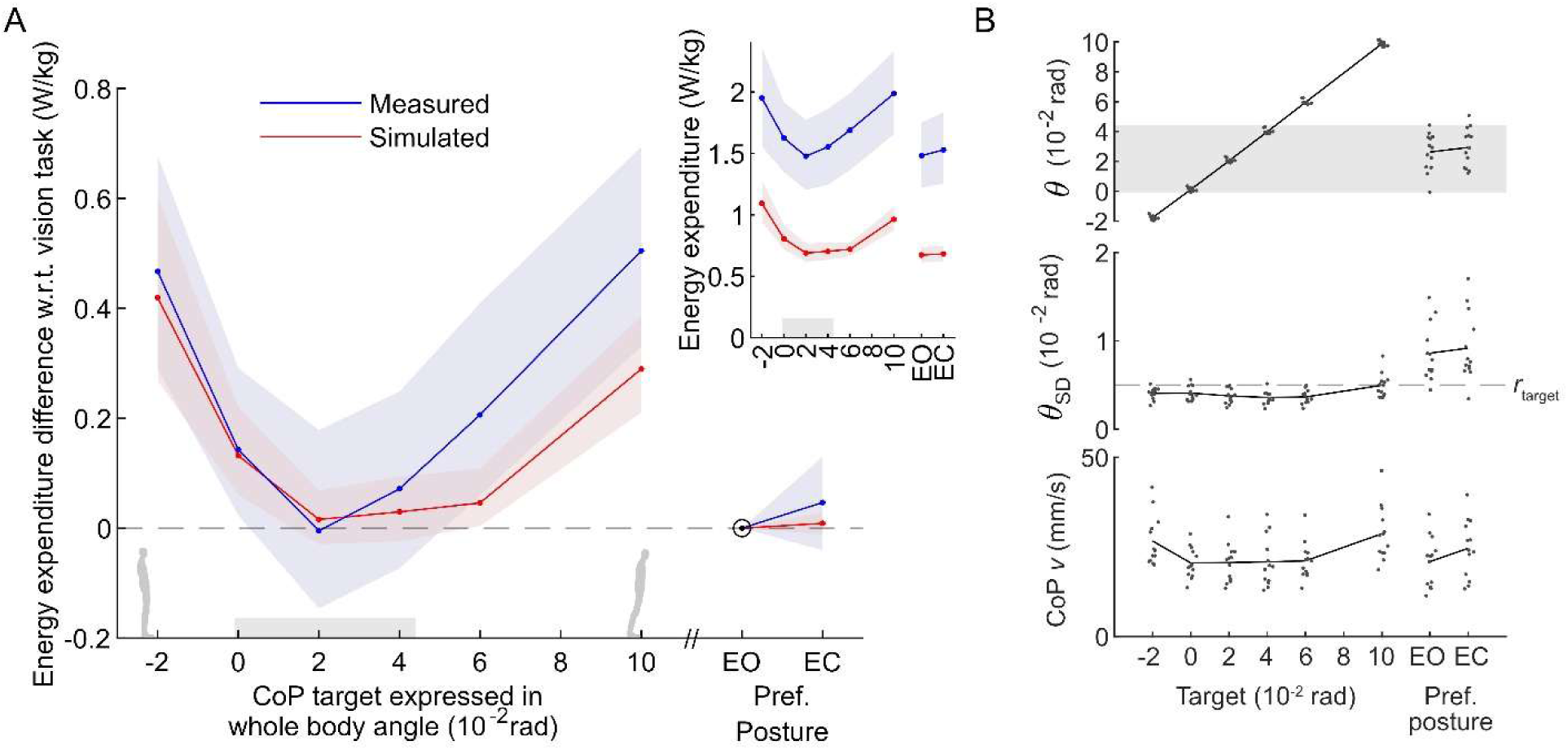
Metabolic energy expenditure of different postures in Experiment 1. A: The means and bootstrapped 95% confidence interval of the measured (blue) and simulated (red) metabolic expenditure at each of the target conditions (−2, 0, 2, 4, 6, and 10 · 10^-2^ rad, equal to −1.15, 0, 1.15, 2.29, 3.44, 5.73°) and the preferred posture with eyes open (EO) and eyes closed (EC). The main panel shows the mass-normalized energy expenditure difference relative to the EO condition (subtraction performed for each participant independently). The inset shows the mass-normalized absolute energy expenditure. Note that the simulated expenditure as computed from the musculoskeletal simulation only considers lower-limb skeletal muscles and omits contributions from any other source, whereas indirect calorimetry measurements represent whole-body energy expenditure. B: Participant means for whole-body angle *θ*, within-trial standard deviation of whole-body angle *θ*_SD_, and average magnitude of the CoP velocity in the forward-backward direction during target and preferred posture trials. The radius of the target is denoted as *r*_target_. The range between the most posterior and anterior preferred postures observed in the EO condition (−0.1 to 4.4 · 10^-2^rad) is marked gray in both panels.

During preferred posture trials, participants self-selected a posture throughout the trial, and most participants (12/13) adopted a slight forward lean in both the eyes open and closed conditions. The range of average preferred postures across participants spanned from −0.1 to 4.4 · 10^-2^ rad (−0.03 to 2.54°) for eyes open and 1.2 to 5.1 · 10^-2^ rad (0.70 to 2.91°) for eyes closed conditions (see Figure 2B). On average, these whole-body lean angles did not differ significantly between the eyes open and closed conditions (EO: 2.6 ± 1.3 · 10^-2^ rad (1.50 ± 0.73°); EC: 2.9 ± 1.3 · 10^-2^ rad (1.68 ± 0.74°), *t*(12) = −2.099, *p* = 0.058). Similarly, there was no significant difference in measured energy expenditure between eyes open and closed standing (EO: 1.48 ± 0.48 W/kg; EC: 1.53 ± 0.54 W/kg; *t*(12) = –1.076, *p* = 0.303; Figure 2A inset).

In the target condition trials, participants controlled a cursor on a screen (see Figure 1A and 1C) that represented their center of pressure (CoP) as measured by the force plates. The energetic cost measured through indirect calorimetry was minimal at the 2 · 10^-2^ rad (1.15°) target and increased in both lean directions. At this target, the energy expenditure was not significantly different from that of preferred posture with eyes open (EO: 1.48 ± 0.48, Target: 1.48 ± 0.52 W/kg, *t*(12) = 0.059, *p*_Holm_ = 0.954, Holm corrected for six comparisons, Figure 2A). Across all postures, the energy expenditure varied significantly with the prescribed target (rmANOVA, *F*(5, 60) = 14.11, *p*_target_ = < 0.001, Figure 2A). To assess how energetic cost increased with respect to the minimum-cost posture, we performed planned post-hoc comparisons of energy expenditure across targets to the 2 · 10^-2^ rad target (1.48 ± 0.52 W/kg) using one-sided *t*-tests. The difference with respect to the 2 · 10^-2^ rad target was significant for all posterior postures (Target −2: 1.95 ± 0.73 W/kg, *t*(12) = −5.42, *p*_Holm_ < 0.001; Target 0: 1.62 ± 0.52 W/kg, *t*(12) = −3.27, *p*_Holm_ = 0.007) and for most more anterior postures (Target 4: 1.55 ± 0.57 W/kg, *t*(12) = −0.87, *p*_Holm_ = 0.201; Target 6: 1.69 ± 0.57 W/kg, *t*(12) = −3.89, *p*_Holm_ = 0.007, Target 10: 1.99 ± 0.62 W/kg, *t*(12) = −10.94, *p*_Holm_ < 0.001). This pattern of increasing energy expenditure at larger anterior and posterior lean angles was accompanied by a small increase in minute volume (total air circulated per minute) at the outer targets (Target −2 vs 0 to 4, Targets 0 to 6 vs 10; see S1 Appendix for statistics) and a slight increase in breath frequency (Target −2 vs 0, Targets 0 to 6 vs 10). The minute volume was not significantly different between the eyes open preferred posture and the minimum-cost target (EO: 10.2 ± 3.4, Target 2: 10.3 ± 3.1 L/min, *t*(12) = −0.102, *p*_Holm_ = 0.920), although the breathing frequency was slightly higher (EO: 14.6 ± 3.70, Target: 18.5 ± 4.9 breaths/min, *t*(12) = −3.201, *p*_Holm_ = 0.016).

To test whether the increase in metabolic cost was symmetric around the minimum-cost target (2 · 10^-2^ rad), we calculated the average rate of change in energy expenditure across adjacent targets in both the anterior and posterior directions from this minimum. The average slope in the posterior direction was −11.80 ± 7.84 W/kg/rad (−0.21 ± 0.14 W/kg/°), whereas the slope in the anterior direction was 6.01 ± 2.57 W/kg/rad (−0.10 ± 0.04 W/kg/°). The absolute value of these slopes differed significantly (*t*(12) = 3.763, *p* = 0.003), suggesting that the energetic cost of posture was asymmetric, increasing at almost twice the rate in the posterior direction. Overall, these results show that energy expenditure during quiet standing is a function of whole-body lean angle, with a clear energetic minimum near a slight forward lean. Importantly, the increase in cost was asymmetric, rising more steeply in the posterior direction.

### Preferred postures are variable but centered around the minimum-cost whole-body lean

Although participants generally preferred postures near the energetic minimum, their whole-body orientation also showed substantial variation both within and across individuals (see Figure 2B). To analyze within-trial variability, we analyzed the center of pressure velocity in the forward-backward direction and the standard deviation of whole-body lean angle. We first compared responses from the preferred posture trials (eyes-open) with the minimum-cost posture (i.e., 2 · 10^-2^ rad), which was also the target nearest to the preferred posture (Figure 2A). The average magnitude of the CoP velocity, which is an indication of the magnitude of postural corrections, did not differ between the preferred posture and the minimum-cost target posture (EO: 21 ± 7, Target 2: 22 ± 5 mm/s, *t*(12) = 0.163, *p* = 0.873, Figure 2B). The within-trial standard deviation of lean angle was roughly double that of the minimum-cost target (EO: 0.92 ± 0.32 · 10^-2^ rad (0.49 ± 0.18°), Target 2: 0.38 ± 0.08 · 10^-2^ rad (0.22 ± 0.05°), *t* = 5.774, *p*_Holm_ < 0.001, Figure 2B), likely because the task to remain within the target constrained the allowed orientations. This suggests that variability in whole-body orientation (i.e., standard deviation of lean angle) was greater during preferred standing than at the minimum-cost target, but this did not translate into increased postural corrections (CoP velocity) or higher energy expenditure.

Next, we compared these same responses across the target trials. While the average CoP velocity differed across the targets (rmANOVA, *F*(2.62, 31.4) = 19.61, *p*_target_ = < 0.001, Greenhouse-Geisser corrected), pairwise comparisons revealed significant differences only between the central postures (0 to 6 · 10^-2^ rad) and the most posterior and anterior leaning postures (−2 and 10 · 10^-2^ rad, see S1 Appendix for statistics). Similarly, the standard deviation of the whole-body angle was also significantly different across postures (rmANOVA, *F*(2.07, 24.87) = 13.22, *p*_target_ = < 0.001, Greenhouse-Geisser corrected); however, pairwise comparisons revealed that only the most anterior target was significantly differed from all others (10 vs −2, 0, 2, 4, and 6 · 10^-2^ rad, see S1 Appendix for statistics). This suggests that minimizing whole-body variability, a common objective in postural control models, may not be a prerequisite for minimizing energy expenditure. Finally, closing the eyes did not change the within-trial variability of whole-body lean (EC: 0.93 ± 0.38 · 10^-2^ rad (0.53 ± 0.22°), *t*(12) = −0.753, *p* = 0.466, Figure 2B), but modestly increased the center of pressure velocity (EO: 20 ± 7; EC: 26 ± 8 mm/s, *t*(12) = −4.385, *p* = 0.001, Figure 2B) compared to preferred posture with eyes open. To identify possible mechanical sources of across-participant variability, we assessed whether individual variability in ankle stiffness could account for differences in preferred posture. When standing is simplified as a single inverted pendulum, the actively generated moment is minimal when the passive moments generated by the tissues around the ankles counteract the gravitational moment acting on the body. We used a published model of ankle stiffness (42) to estimate a distribution of whole-body lean angles where the passive ankle moment is in equilibrium with the gravitational moment of a body with a height equivalent to our group average (see Materials and Methods). Using a Monte Carlo analysis, we found that the simulated equilibrium angles formed a left-skewed normal distribution with a median of 2.2 · 10^-2^ rad and an interquartile range of 0.9 [1.8, 2.7] · 10^-2^ rad (data reported in non-parametric metrics to account for skewness of the distribution). For comparison, the preferred postures measured in Experiment 1 had a median of 2.9 · 10^-2^ rad and an IQR of 2.0 [1.6, 3.6] · 10^-2^ rad. This suggests that individual ankle mechanics may partly explain between-participant differences in preferred posture, but also shows that participants generally adopted more forward-leaning postures than predicted by the passive equilibrium angles alone.

### Musculoskeletal model simulations predict the metabolic cost in posture

To better understand the physiological basis of postural energy expenditure, we used a musculoskeletal model in OpenSim (12, 13) to estimate muscle-level energetic cost during standing (Figure 1B). Muscle energy expenditure was computed for 80 muscles using a physiological model (30) that accounted for the muscle’s activation, mass, fiber composition, and (tendon) stiffness. The basal metabolic cost was subtracted from the measured and simulated costs separately, allowing for a direct comparison of the cost increases (Figure 2A). Similar to our experimental data, the energy expenditure in our simulations was lowest at 2 · 10^-2^ rad and varied significantly with the target lean angle (rmANOVA, *F*(1.62, 19.4) = 15.63, *p*_target_ = < 0.001, Greenhouse-Geisser corrected), increasing at both posterior and anterior orientations. Relative to the minimum-cost posture (2 · 10^-2^ rad: 0.69 ± 0.13 W/kg), post-hoc comparisons revealed that energy expenditure increased significantly at all posterior postures (Target −2: 1.09 ± 0.31 W/kg, *t*(12) = −5.03, *p*_Holm_ < 0.001; Target 0: 0.80 ± 0.19 W/kg, *t*(12) = −3.01, *p*_Holm_ = 0.016, Figure 2A), and at the most anterior posture (Target 4: 0.70 ± 0.13 W/kg, *t*(12) = −0.68, *p*_Holm_ = 0.318; Target 6: 0.72 ± 0.10 W/kg, *t*(12) = −1.04, *p*_Holm_ = 0.318; Target 10: 0.96 ± 0.17 W/kg, *t*(12) = −5.45, *p*_Holm_ < 0.001). Overall, simulations captured well the trends in measured energy expenditure (see Figure 2A).

To validate the simulated muscle activations and explore posture-dependent muscle recruitment, we compared experimental EMG recordings with simulated activations for eight major postural muscles (Figure 3). Overall, the simulations captured the qualitative trends in EMG across postures, although absolute magnitudes likely differed due to normalization (43) (see Methods). Muscles responsible for dorsiflexion (tibialis anterior) and knee extension (rectus femoris, vastus medialis) were highly active in backward postures, with activation declining as the body leaned forward. In contrast, forward postures increased reliance on ankle plantarflexors (soleus, gastrocnemius) and hip extensors (semitendinosus, semimembranosus). To better understand the different physiological contributions to energy expenditure, we decomposed the total metabolic cost from the simulations into activation maintenance, fiber shortening rate, and mechanical work (30). The activation maintenance reflects the metabolic losses of sustaining a constant, isometric muscle force, including processes such as calcium cycling and cross-bridge turnover (30). The shortening rate accounts for additional losses occurring when the muscle fibers change length, while mechanical work represents the muscle’s idealized external work (30). During preferred posture trials, nearly all energy expenditure was attributed to activation maintenance costs (99.4 ± 0.6%), with only a minimal contribution from the fiber shortening rate (0.6 ± 0.6 %) and none from mechanical work. This confirms that static standing primarily expends energy through sustained low-level muscle activation, rather than dynamic contraction or external work.

**Figure 3:**
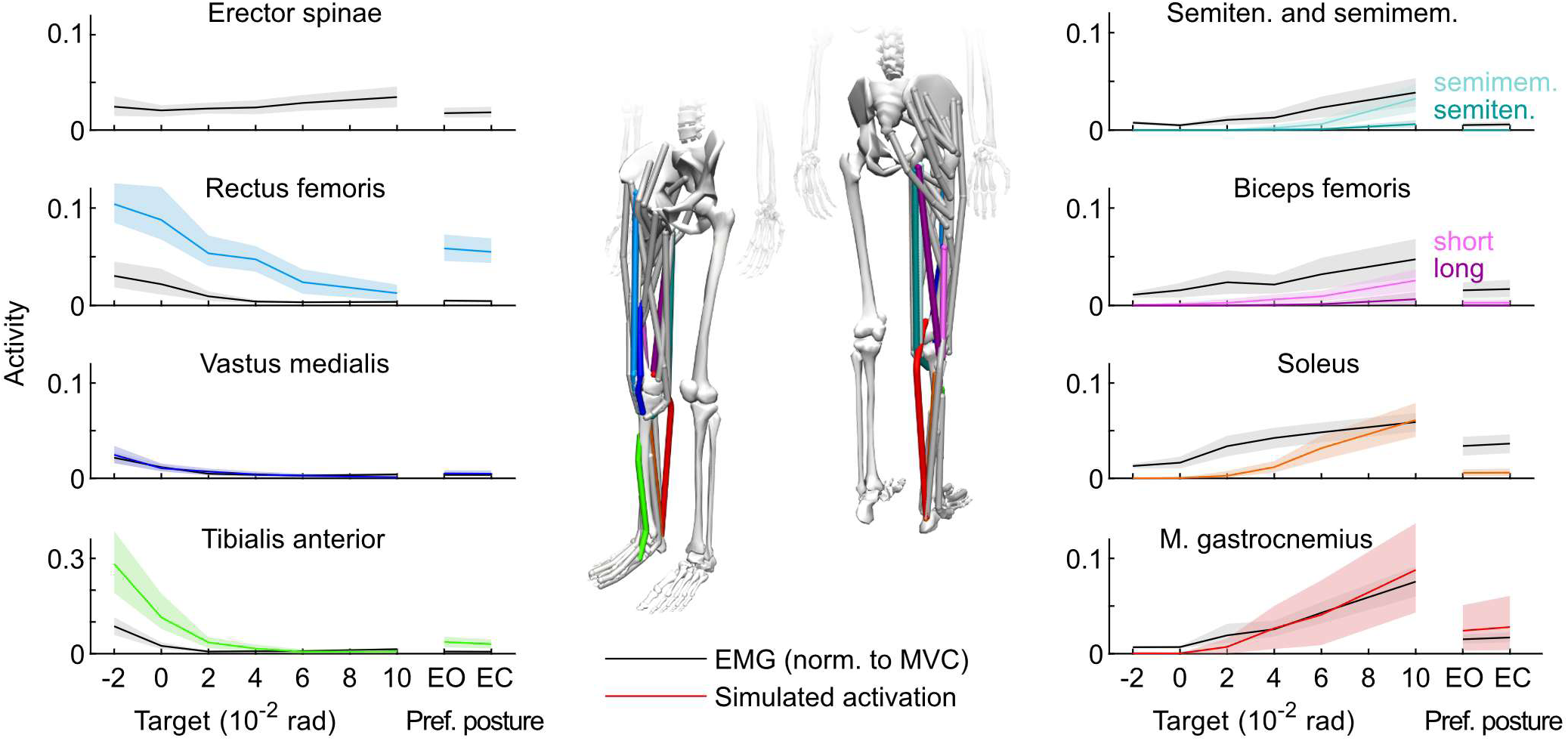
Measured (black) and simulated (colored) activity of major postural muscles on the right side of the body during Experiment 1. Solid lines indicate the mean activity across participants, and the shaded areas represent the bootstrapped 95% confidence interval of the mean. EMG data were collected for the erector spinae (not present in the musculoskeletal model), rectus femoris, vastus medialis, tibialis anterior, semitendinosus, biceps femoris (long head), soleus, and medial gastrocnemius muscles. The activation of the semimembranosus is also shown, as it shares a similar function with the semitendinosus. Electromyography data were normalized to the largest value observed during maximal voluntary contraction (see Methods), and simulated muscle activations were inherently constrained between zero and one.

Interestingly, the proportion of energy expended by muscles consisting mostly of more energy-intensive fast-twitch muscle fibers (type II) also varied with lean angle. At the most posterior target (Target −2), 70.0 ± 9.4% of the simulated energy expenditure came from muscles with relatively high (> 52%) fast-twitch fibers, such as the rectus femoris and vastus lateralis (29). In contrast, at the most anterior target (Target 10), only 42.6 ± 13.8% of expended energy came from such muscles, with the remainder of the energy being consumed by muscles consisting primarily of slow-twitch fibers (type I), such as the soleus (29). This result suggests that posterior leaning may recruit less metabolically efficient muscles for prolonged activation maintenance, contributing to the higher energetic cost observed when leaning in that direction.

### Preferred quiet standing posture is not affected by the metabolic cost of gait initiation

So far, we observed that energy expenditure during static standing was minimal when participants adopted a whole-body orientation at 2 · 10^-2^ rad anteriorly (Figure 2), closely matching their preferred posture when standing freely. However, the optimization of posture may reflect more than just minimizing the cost of static standing; it may also be shaped by the need to initiate movement quickly, efficiently, and frequently (39, 44). To explore this proposition, Experiment 2 investigated how the energetic cost and time required to initiate gait vary with posture and whether participants adjust their static posture in anticipation of the movement direction. Participants (*N* = 20) stood on a force plate at a target or at their preferred posture and initiated gait when prompted to start walking in a forward or backward direction (Figure 1F-G, see Materials and Methods). Participants were explicitly informed as to which set of trials (Forward, Backward, Random) they were performing, and could therefore anticipate the probability of the walking direction but not the timing of the walking prompt.

The lean angle assumed by participants during standing before gait initiation did not differ significantly between sets where the direction was randomized or known (Random: 0.018 ± 0.018 rad (1.01 ± 1.02°); Forward: 0.017 ± 0.018 rad (1.00 ± 1.01°), *t*(19) = 0.094, *p* = 0.926; Backward: 0.017 ± 0.019 rad (0.96 ± 1.06°), *t*(19) = 0.734, *p* = 0.460; Figure 4B). Similarly, the time required to reach steady-state walking (i.e., when the center of mass velocity reached 90% of their average maximum velocity for that direction) was not significantly different when the direction was randomized or known in both the forward (Random: 1.62 ± 0.21 s vs. Forward: 1.56 ± 0.23 s; *t*(19) = 1.455, *p* = 0.162, Figure 4B) and backward directions (Random: 1.25 ± 0.15 s vs. Backward: 1.26 ± 0.20 s; *t*(19) = −0.172, *p* = 0.865). Nevertheless, the time to reach steady state walking was shorter during backward as compared to forward walking initiation (*t*(19) = −6.051, *p* < 0.001, Figure 4B), while the maximal absolute velocity in backward walking was significantly lower (Forward: 1.13 ± 0.17 m/s vs. Backward: 0.92 ± 0.14 m/s; *t*(19) = −11.3, *p* < 0.001, Figure 4B).

**Figure 4:**
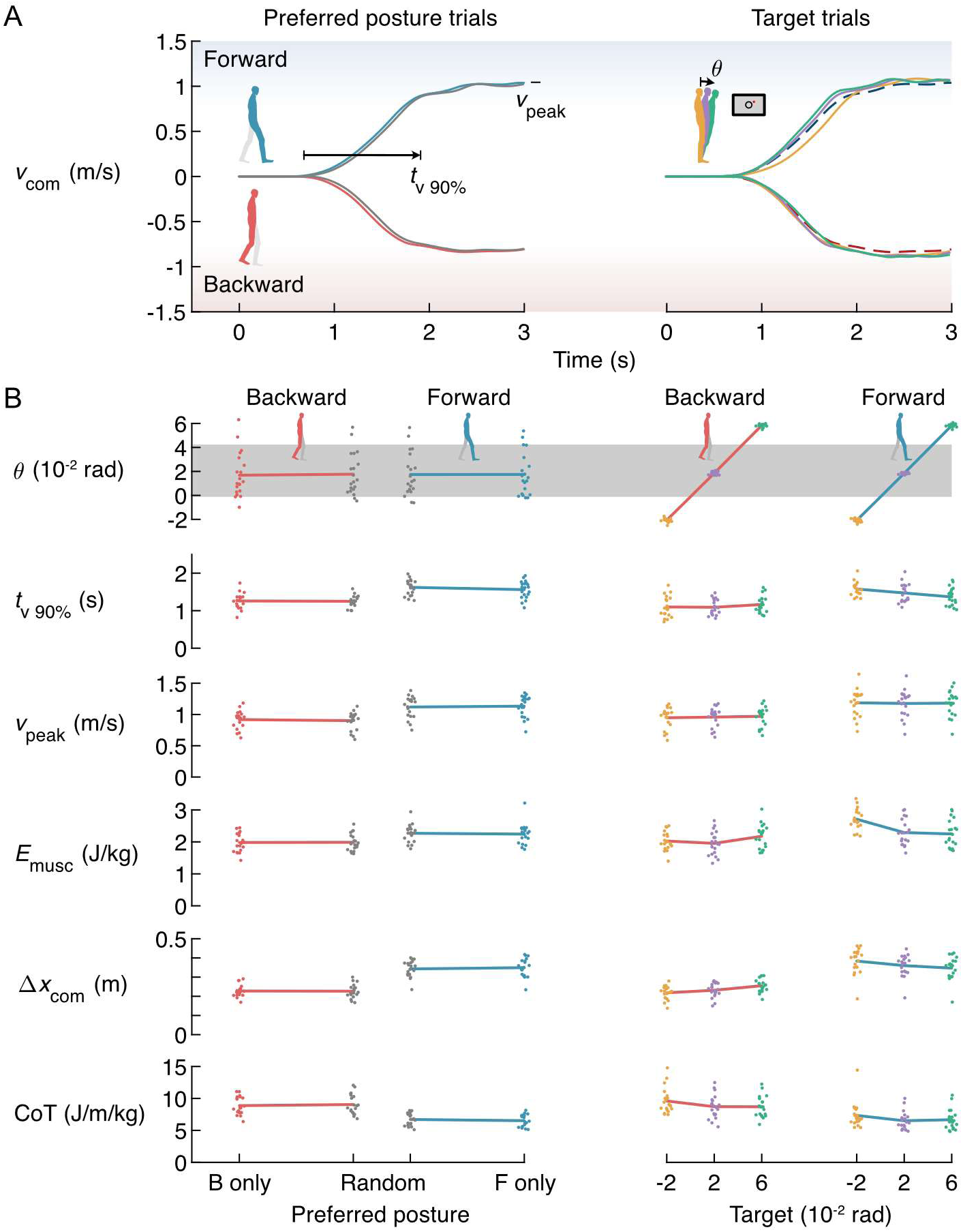
Gait initiation results for preferred posture and target trials of Experiment 2. In preferred posture trials, the participant could self-select a posture during the period before the cue. In target trials, participants were prescribed a lean angle using the CoP target. A: Mean forward velocity of the center of mass for all participants (N=20) grouped by trial. Blue and red lines correspond to the Forward- and Backward-only block, where participants were aware that they would only initiate gait in the corresponding direction (repeated as dashed lines in the right figure for comparison). The gray lines (lagging slightly behind the blue and red lines) show the initiation in the Random direction block, where the direction was unpredictable. The colored lines (yellow, purple, green) correspond to the three prescribed targets (−2, 2, and 6 · 10^-2^ rad, −1.15, 1.15, 3.44°, respectively). B: Mean results of gait initiation metrics. The gray band indicates the range of preferred postures for prolonged standing as observed in Experiment 1. The reported metrics are the whole-body lean angle theta, time to reach 90% of steady-state velocity with respect to the start of movement, peak forward velocity (absolute value), computed energy expenditure by all muscles from the presentation of the prompt at time 0 to the first foot strike, change in center of mass location between time 0 and first foot strike, and the cost of transport (CoT) over that period, equal to *E*_musc_ / Δ*x*_com_.

Our simulations of Experiment 1 showed that activation maintenance costs drive energy expenditure during quiet standing. However, many idealized representations of gait initiation only consider kinetic and potential energy (45–48), overlooking the cost of losses from muscle activation and likely underestimating total energy expenditure. To address this, we further used the musculoskeletal model to estimate energy expended by the lower-limb muscles during gait initiation from the presentation of the cue to the first foot strike (heel or forefoot, Figure 1B). The cost of gait initiation was quantified using the cost of transport (CoT) of the center of mass between the presentation of the prompt (i.e., during standing) and the first foot strike, and was normalized to the participants’ body weight. The CoT was computed as the energy expended by the simulated muscles up to the first foot strike (*E*_musc_) divided by the change in center of mass location in the forward direction (Δ*x*_CoM_) as determined from inverse kinematics (Figure 4B). During the sets with known walking direction (Forward and Backward sets), the CoT was significantly lower for forward (6.52 ± 0.95 J/m/kg, Figure 4B) compared to backward initiation (8.87 ± 1.38 J/m/kg, *t*(19) = −9.283, *p* < 0.001). When the walking direction was randomly selected, the CoT was slightly higher as compared to the known walking direction in the forward (6.71 ± 0.94 J/m/kg, *t*(19) = 2.380, *p* = 0.028) but not the backward initiation (9.05 ± 1.55 J/m/kg, *t*(19) = 0.638, *p* = 0.531). Together, these results indicate that knowledge of the upcoming gait initiation direction did not alter participants’ static standing posture, even though we observed a marginal improvement in estimated energy expenditure for forward initiation.

### Static lean orientation only minimally affects the energy and time expended for gait initiation

In the second block of Experiment 2, we used the target trials to test whether initiating gait from a more anterior or posterior leaning whole-body angle improves the time required and/or the energetic cost to initiate gait. Participants stood at one of three prescribed whole-body orientations (−2, 2, and 6 · 10^-2^ rad, Figure 1D) and initiated walking in either the forward or backward direction five times each in random order following a prompt (Figure 1F).

The time to reach steady-state velocity varied significantly with lean angle during forward gait initiation (−2: 1.58 ± 0.18 s, 2: 1.47 ± 0.25 s, 6: 1.37 ± 0.24 s, rmANOVA, *F*(2, 38) = 7.998, *p*_target_ = 0.001, Figure 4B), with post-hocs showing that participants reached steady-state walking faster in the forward-leaning posture as compared to the backward-leaning (*t* = 4.00, *p*_Holm_ < 0.001). However, comparisons of the outer two targets with the central target were not significant (−2 vs 2: *t* = 1.97, *p*_Holm_ = 0.099; 2 vs 6: *t* = 2.03, *p*_Holm_ = 0.099), suggesting that postural adjustments beyond the central posture did not yield significantly faster gait initiation times. In contrast, during backward gait initiations, lean angle had no significant effect on the time to reach steady state velocity (−2: 1.10 ± 0.27 s, 2: 1.09 ± 0.20 s, 6: 1.16 ± 0.22 s, rmANOVA, *F*(2, 38) = 1.797, *p*_target_ = 0.180). The peak center of mass velocity did not differ across targets for forward initiation (−2: 1.18 ± 0.21 m/s, 2: 1.18 ± 0.21 m/s, 6: 1.18 ± 0.20 m/s, rmANOVA, *F*(2, 38) = 0.79, *p*_target_ = 0.425, Greenhouse-Geisser corrected), but was marginally higher (0.02 m/s) when initiating backward walking from a forward-leaning posture versus a backward-leaning posture (−2: 0.95 ± 0.17 m/s, 2: 0.96 ± 0.15 m/s, 6: 0.97 ± 0.15 m/s, rmANOVA, *F*(2, 38) = 5.19, *p*_target_ = 0.010; post-hoc −2 vs 6: *p*_Holm_ = 0.008). Overall, these results suggest that the time required to initiate gait in either the forward or backward direction can be improved slightly by adjusting whole-body orientation in quiet standing.

The CoT differed with lean angle for both forward (Target −2: 7.34 ± 1.93 J/m/kg, Target 2: 6.51 ± 1.39 J/m/kg, Target 6: 6.67 ± 1.57 J/m/kg, *F*(1.24, 23.64) = 7.68, *p*_target_ = 0.007, Greenhouse-Geisser corrected) and backward gait initiations (Target −2: 9.60 ± 1.97 J/m/kg, Target 2: 8.86 ± 1.65 J/m/kg, Target 6: 8.69 ± 1.63 J/m/kg, *F*(1.37, 26.06) = 5.044, *p*_target_ = 0.024, Greenhouse-Geisser corrected). Interestingly, post-hoc tests revealed that for both initiation directions, the CoT was higher when leaning at the posterior target as compared to the central target (Forward −2 vs 2: *p*_Holm_ = 0.002, Backward −2 vs 2: *p*_Holm_ = 0.040), contrary to the hypothesis that leaning in the posterior direction would facilitate backward initiation. Conversely, the CoT did not differ between the anterior and central target for both initiation directions (Forward: 2 vs 6: *p*_Holm_ = 0.470, Backward 2 vs 6: *p*_Holm_ = 0.578), suggesting that maintaining a posture close to 2 · 10^-2^ rad was sufficient to optimize the CoT of gait initiation regardless of direction.

To better understand what drives the difference in CoT between anterior and posterior leaning postures, we examined both the total muscle energy expended (*E*_musc_) and the CoM displacement individually. The total muscle energy expended (as estimated from our simulations) between the cue to walk and first foot strike decreased significantly when leaning in the direction of movement for both forward (Target −2: 2.70 ± 0.32 J/kg, Target 2: 2.29 ± 0.40 J/kg, Target 6: 2.25 ± 0.42 J/kg, *F*(1.37, 25.94) = 39.902, *p*_target_ < 0.001, Greenhouse-Geisser corrected) and backward initiations (Target −2: 2.03 ± 0.29 J/kg, Target 2: 1.99 ± 0.34 J/kg, Target 6: 2.18 ± 0.37 J/kg, *F*(1.62, 30.72) = 10.563, *p*_target_ < 0.001, Greenhouse-Geisser corrected). Differences in CoT between the central target and the target aligned with the direction of movement were not significant (Forward, Target 2 vs 6: *t*(19) = 0.749, *p*_Holm_ = 0.459; Backward, Target −2 vs 2: *t*(19) = 0.831, *p*_Holm_ = 0.411), whereas initiating gait while leaning opposite to the movement direction incurred a higher energetic cost than both other targets (*p*_Holm_ < 0.003 for all comparisons). Furthermore, the center of mass displacement between the starting position and the first foot strike also decreased when leaning further in the direction of movement for both forward (Target −2: 0.38 ± 0.07 m, Target 2: 0.36 ± 0.0.06 m, Target 6: 0.35 ± 0.06 m, *F*(1.46, 27.73) = 16.66, *p*_target_ < 0.001, Greenhouse-Geisser corrected) and backward trials (Target −2: 0.22 ± 0.03 m,

Target 2: 0.23 ± 0.04 m, Target 6: 0.25 ± 0.04 m, *F*(1.89, 35.87) = 28.28, *p*_target_ < 0.001, Greenhouse-Geisser corrected). Because CoT is energy expenditure per distance traveled, these combined reductions in both energy and displacement cancelled each other out, suggesting that leaning further in the direction of movement does not necessarily improve energetic efficiency (i.e., CoT). Instead, maintaining a posture close to 2 · 10^-2^ rad appears sufficient to minimize the CoT during gait initiation, while leaning further in the direction of travel offers little to no reduction in energetic cost or time spent to reach steady-state walking.

## Discussion

In this study, we examined (i) how metabolic cost varies with standing posture, (ii) how the cost of gait initiation depends on posture, and (iii) whether humans adopt postures that minimize cost during quiet standing and in preparation for walking. In Experiment 1, we quantified the metabolic cost of posture across a range of natural standing postures, which increased systematically but asymmetrically with lean angle and was lowest at a slight forward lean (2 · 10^-2^ rad; 1.15°). Participants’ preferred postures that were close to this energetic minimum (2.6 ± 1.3 · 10^-2^ rad; 1.50 ± 0.73°). Musculoskeletal simulations matched the experimentally measured energy expenditure, validating that these models can be used to predict energy expenditure during standing. In Experiment 2, participants continued to prefer a forward-leaning orientation and did not adjust their posture for gait initiation, even when the upcoming gait direction was known. Consistent with this, simulations of gait initiations showed that the energy cost of gait initiation was also near optimal at a slight forward lean (2 · 10^-2^ rad), regardless of the direction of gait initiation. Taken together, our results indicate that participants maintained a preferred posture close to the minimum-cost orientation for both static standing and gait initiation, supporting the idea that we minimize energy expenditure during quiet standing. However, substantial within- and between-participant variability in preferred posture suggests that the minimization of energy expenditure does not require strictly matching a single energy-optimal reference posture. These findings provide insight into how energy expenditure may underly our everyday movement control and challenge the common modeling assumption that postural costs can be described accurately using a symmetric error from a single reference posture (e.g., vertical or static equilibrium with ankle stiffness).

### Preferred human posture aligns with the energetic minimum

The first aim of our study was to determine how energy expenditure varies with standing posture. The measured energy expenditure was lowest at a target lean angle of 2 · 10^-2^ rad in the anterior direction, closely aligning with the average preferred posture of 2.6 ± 1.3 · 10^-2^ rad when participants had their eyes open. The alignment of preferred posture with the energetic minimum suggests that participants naturally adopted an energetically efficient posture during quiet standing. We also found that removing visual feedback did not significantly affect energy expenditure or lean angle, although center of pressure (CoP) velocity increased slightly. This indicates that while visual feedback aids postural control (for review, see 27, 49, 50), its absence does not substantially alter the selection or energetic cost of the preferred posture.

Our results further revealed that the increase in energy expenditure was not symmetrical (Figure 2A), increasing at about twice the rate during posterior lean as compared to anterior lean. This directional asymmetry in energetic cost likely reflects underlying structural and physiological asymmetries in the human body, which may have evolved under selection pressures for efficient (forward) locomotion (51, 52) and shape the energetic landscape of standing balance. For instance, the relatively short calcaneus, while advantageous for energy-efficient running (53), limits posterior stability because it is much shorter than the forefoot and therefore requires faster CoP corrections (Figure 2B) to prevent backward falls. Similarly, passive ankle stiffness increases with forward lean (42), reducing the proportion of active muscle contributions needed to generate stabilizing torques. Lastly, backward-leaning postures engage muscles dominated by slow-twitch fibers, which are less metabolically efficient, whereas forward lean relies on fast-twitch fibers better suited for economical tonic activation (30, 52). Together, these biomechanical and muscular properties make it less costly to allow greater excursions in the anterior direction than in the posterior, helping to explain why humans typically adopt forward-leaning stance.

Additional non-metabolic factors may also drive our preference for a slight forward lean. Sinha and Maki (54) proposed that this orientation simplifies control by only relying on tonic plantarflexor activity, while also increasing the gain of short-latency reflexes through increased tonic activity (55). Second, a slight forward lean places the center of mass closer to the center of our asymmetric base of support, which could align with optimizing our stability boundaries (56, 57). Lastly, this preference may also prepare us to initiate movement (44), although our direct examination of this possibility in Experiment 2 showed that participants remained near their energy-optimal stance regardless of the expected walking direction (see below). Even so, other influences not assessed here may also contribute to posture selection, including the need to stabilize the head (58), minimize cognitive demands of control (59, 60), or allow exploratory variability (61).

### Costs of gait initiation do not drive quiet standing posture

The second aim of this study was to determine how quiet standing posture affects the cost of gait initiation and whether this cost influences our choice of standing posture. We found that leaning further in the intended direction of movement provided no significant energetic benefit or reduction in time spent initiating gait when compared to the minimum-cost static posture at 2 · 10^-2^ rad. For forward walking, leaning further forward as compared to a central posture (6 vs 2 · 10^-2^ rad) only reduced the time to reach steady-state velocity by 0.11 ± 0.23 seconds and reduced the CoT of the first step by 0.17 ± 0.49 J/m/kg, consistent with prior reports that posture has little impact on the characteristics of forward gait initiation (40). For context, humans adjust their steady-state walking speed to minimize CoT differences in the order of ∼0.42 J/m/kg (0.1 kcal/km/kg), and continuously expend around 3.8 J/m/kg (6). Even initiating gait from the least efficient posture tested (−2 · 10^-2^ rad), which increased energy expenditure by 0.42 ± 0.27 J/kg relative to the minimum-cost posture, would raise total expenditure over 10 m of walking by only ∼1%. Such marginal differences indicate that the energetic penalties of starting from a suboptimal stance are too small to drive systematic adjustments. Consistent with this, participants maintained their preferred static posture even when the direction of gait initiation was predictable. Together, these findings suggest that while posture can modulate the cost and timing of gait initiation at larger lean angles (i.e., an observed significant main effect), the influence is negligible relative to the continuous cost of walking and standing, and thus unlikely to shape postural choice under typical everyday conditions.

Instead, it may be possible that time and task constraints have a stronger influence on postural selection for gait initiation. For example, sprinters adopt an energetically costly crouched start to maximize acceleration, whereas marathon runners begin upright to conserve energy. Clearly, urgency can drive changes in posture when the task demands it (44). In our experiment, participants had no incentive to minimize the time spent initiating gait, as the speed of their initiation did not hinder their ability to perform the task effectively. We hypothesize that stricter time demands, such as reaching the end of the walkway as quickly as possible, would have prioritized acceleration at the expense of energetic efficiency (45). Future work could assess how humans optimize energy expenditure using postural adjustments in gait initiation tasks with greater time pressure.

### Variability and flexibility in postural control

In both experiments, participants naturally converged on an energetically efficient posture when no target was prescribed, yet also exhibited substantial variability both across participants and within trials (Figure 2 and 4). A Monte Carlo analysis of a passive ankle stiffness model showed that differences in participants’ stiffnesses could explain only part of this variability. The simulated equilibrium angles (median 2.2 · 10^-2^ rad, IQR = 0.9 [1.8, 2.7] · 10^-2^ rad) – where active contributions to the ankle moment are minimal – were slightly more posterior than the actual postures adopted in Experiment 1 (median 2.9 · 10^-2^ rad, IQR = 2.0 [1.6, 3.6] · 10^-2^ rad). This suggests that a participant’s preferred posture is not solely the result of minimizing active contributions to the ankle moment, consistent with earlier work showing that humans typically adopt a more forward-leaning stance than predicted by inverted pendulum models (23, 62).

Both metrics for within-trial postural variability (i.e., standard deviation of lean angle and CoP velocity, Figure 2B) did not differ across central targets (0 to 6 · 10^-2^ rad). This aligns with recent work showing that variability in postural control signals (i.e., ankle torque) remains relatively constant within this range and rises beyond it, likely due to increased motor noise at higher force demands (63). Furthermore, when standing at preferred postures, the within-trial standard deviation of lean angle roughly doubled relative to the minimum-cost target (Figure 2B), consistent with larger variability and low-frequency drifts previously reported in unconstrained posture (64, 65). This larger dispersion of orientations did not coincide with higher CoP velocity or metabolic cost (Figure 2), suggesting that variability in a low-cost region can be permitted without energetic penalty. Indeed, optimal control models of balance show that strictly limiting center of mass or center of pressure variability can actually hinder energy minimization (28, 37), though importantly, this does not restrict energetic costs from rising when variability is increased through more challenging experimental conditions (36). Intermittent balance control models (66–68) formalize this tradeoff of effort and variability by permitting postural sway until a stability threshold is exceeded, thereby reducing energetic demands compared to continuous control (69). Overall, minimizing variability does not appear to be a prerequisite for minimizing energy expenditure, suggesting that efficient postural control does not require constraining posture at a single energetic minimum. The implications of this observation on modeling methods are discussed further below.

### Towards improved models of cost in posture

Postural control is commonly simplified as a continuous minimization of deviations from a reference lean angle of an inverted pendulum (21–24), resulting in a symmetric linear or quadratic cost function with a single optimum in the center. However, our experimental data show that energy expenditure increases at almost twice the rate with backward lean than with forward lean, which is not captured by these cost functions. To improve the predictive and exploratory power of inverted pendulum models, we recommend penalizing posterior-leaning postures more strongly than anterior-leaning postures to account for the asymmetries observed in energy expenditure. Future work could evaluate whether multi-objective control models (e.g., a linear quadratic regulator minimizing control action and kinematic state) and intermittent control (66–68) can be improved by incorporating asymmetric weights to control actions in the anterior and posterior directions, which may better represent the cost of accelerating the body in these directions. We hypothesize that this adjustment may explain the human tendency to maintain an anterior-leaning posture during quiet standing (62) as a precaution to avoid the steeper cost gradient associated with a posterior lean.

Alternatively, musculoskeletal simulations may offer a more physiologically grounded approach to represent the energetic costs of posture. Such models have been applied to simulate postural control (25, 26, 70, 71) and estimate the energetic cost of postural responses (72). Our study extends this work by validating that musculoskeletal inverse simulations can reproduce the differences in measured energetic cost of standing, including the asymmetric increase for posterior leaning angles. However, the model underestimated the increase in energetic cost for anterior leaning postures (Figure 2A). One likely explanation is its high passive ankle stiffness: for a participant with parameters close to the unscaled model (male, 1.77 cm, 80 kg), the summed tendon force of the inactive soleus and gastrocnemius (medialis and lateralis) muscles generated 400 N at an ankle angle of 5 degrees, resulting in a moment of 17 Nm. By comparison, Moseley, Crosbie (42) reported a passive ankle moment of only ∼10 Nm at this angle. A second possibility is that contributions from the upper body that are not captured in the model, such as muscles in the arms, neck and back, play an increasing role in energy expenditure for anterior postures. The measured EMG activity of the erector spinae muscle (Figure 3) showed an increase of activity with more anterior postures. Future work could also investigate how individual (sex) differences in morphology, weight distribution, passive ankle stiffness, and muscle strength (73, 74) affect optimal posture.

### Conclusion

This study highlights that human postural control is closely tied to metabolic energy expenditure. By combining indirect calorimetric measurements and musculoskeletal simulations, we quantified the energetic cost of quiet standing and gait initiation over a range of natural standing postures. Metabolic cost was minimal at a slight forward lean and rose asymmetrically with lean angles, at approximately twice the rate in the backward direction. Participants maintained their preferred posture close to this minimum-cost orientation during static standing. Moreover, gait initiation was not a factor for the preference of quiet standing posture, as participants in our second experiment did not adjust their quiet standing posture to the known direction of upcoming gait initiation. Simulations confirmed that preparing for gait initiation by maintaining a whole-body orientation further in the movement direction carried only marginal energetic and temporal benefits, suggesting that maintaining a slight anterior-leaning posture is sufficient to optimize the energetic costs of both standing and gait initiation. However, substantial within- and between-participant variability in preferred posture suggests that the minimization of energy expenditure does not require strictly matching a single optimal reference posture. These findings provide insight into how energy expenditure drives our everyday movement control and challenge the common modeling assumption that postural costs can be described accurately using a symmetric error from a single reference posture. By integrating empirical and simulation-based approaches, our work offers a broader perspective on the cost functions that define optimal postural control and highlights opportunities for their improvement.

## Materials and Methods

This study aimed to determine the energy expenditure required for maintaining a static posture and initiating gait. We performed two experiments: (1) a metabolic assessment of static standing at six prescribed target positions and at preferred postures with eyes open and closed, and (2) a walking initiation experiment where participants stood ready and started walking when presented with a cue.

### Participants

A total of 33 healthy adult participants aged 18-50 with no history of balance, neurological or musculoskeletal disorders were recruited for this study (Experiment 1: 8 male, 5 female, age = 24.2 ± 2.1 years, height = 175.3 ± 7.3 cm, weight = 72.4 ± 9.7 kg; Experiment 2: 9 male, 11 female, age = 24.1 ± 3.3 years, height = 178.1 ± 11.1 cm, weight = 70.0 ± 10.5 kg). All experiments were conducted in accordance with the Declaration of Helsinki and under the approval of the Erasmus Medical Center Medical Ethics Review Committee. Participants received the information letter of the experiment at least 24 hours before participating. Before the experiment was conducted, participants provided written informed consent for their participation, and all participants provided the optional permission to publish the coded data collected during the experiment under a Creative Commons Attribution (CC BY) license.

### Methods and protocol

#### Experiment 1: Static standing protocol

The aim of Experiment 1 was to measure the metabolic cost of anterior and posterior postures and to determine whether this cost could be predicted from motion data using a musculoskeletal model of human standing. A motion capture system (Qualisys AB, Göteborg, Sweden), consisting of 12 Oqus 700 cameras, was used to track the locations of 43 passive motion capture markers placed on the participant at 100 Hz. Marker placement followed the IOR gait marker set (used by (75)) with the addition of four markers on the head, two on each arm, and the omission of the T7 marker due to obstruction by the back-worn metabolic analyzer (See S1 Appendix for list of markers). Ground reaction forces on both feet were measured at 500 Hz using two force plates (9260AA6, Kistler Group, Winterthur, Switzerland), which were zeroed before each set of measurements (four sets per participant) to account for potential signal drift.

Energy expenditure was measured throughout the experiment using a breath-by-breath indirect calorimetry metabolic analyzer (K5 9260AA6, COSMED, Italy). Before each session, the device was calibrated following the manufacturer’s instructions using a certified gas mixture (16% O_2_, 5% CO_2_), a CO_2_ scrubber (C04408-01-07, COSMED, Italy), and a 3 L calibration syringe (C00600-01-11, Hans Rudolph, United States). The analyzer recorded minute ventilation (VE) as well as the O_2_ and CO_2_ concentrations in both inhaled and exhaled air, and reported energy expenditure using the Brockway equation (76). Caloric and caffeine intake are known to affect metabolic assessments (77), so participants were asked not to consume caffeine or large meals 3 hours prior to the experiment’s start time. In addition to this measurement, the energy expenditure was also computed using musculoskeletal simulations (see Analysis).

Electromyography (EMG) data were collected from 8 muscles on the right side of the body, including the medial gastrocnemius, soleus, tibialis anterior, vastus medialis, rectus femoris, biceps femoris, semitendinosus, and erector spinae. The ground electrode was connected to the right ankle on the lateral malleolus, and the SENIAM 8 guidelines (78) were used for electrode placement. Signals were amplified with a gain of 20 and recorded at 2000 Hz using an ExG amplifier (Porti7-8b8at, TMSi, Netherlands). The amplifier was connected to a portable computer (Windows 10, USB 3.0, i7-1195G7, 16 GB RAM, MSI, Taiwan) using a USB connector to start, stop, monitor, and store the EMG data using a modified version of the TMSi MATLAB interface. The measurements were normalized to the maximal observed activation during exercises described by Uhlrich et al. (79, 80) to evoke a maximal voluntary contraction. Briefly, the exercises included prone knee flexion, supine ankle dorsiflexion, hip flexion kicks, and maximum height jumps. A prone lumbar extension exercise without arm support was added to maximize activation of the erector spinae. A synchronization signal was sent from the “sync out” port on the first Oqus camera to the trigger port of the EMG amplifier at the start of each measurement, allowing for the synchronization of the two measurements. The calorimetry measurement was synchronized by manually reading the time on the device at the start of each measurement.

At the start of the experiment, participants were shown the equipment and explained the protocol before signing the informed consent. A short survey was used to collect basic participant information, including age (24.2 ± 2.1 years), sex (8 male, 5 female), dominant hand (12/13 right), and dominant leg (12/13 right). General physical activity was assessed using the short International Physical Activity Questionnaire (Aug 2002 English version) to determine a three-level scale (75, 81). Almost all participants (11/13) were rated at the high activity level, while the remaining participants were rated at the middle level. This distribution was insufficient for making informative comparisons between scores. Next, basic anthropometric measurements were collected, including body length (175.3 ± 7.0 cm), ankle height at the lateral malleolus (6.82 ± 0.6 cm), and hip width at the greater trochanter (34.2 ± 2.1 cm). Body weight (72.4 ± 9.7 kg) was calculated by summing the vertical forces on the force plates and subtracting the weight of the metabolic analyzer (900 g). The foot placement on the force plate was standardized by positioning the centers of the ankle joints at hip width, estimated as 0.532 times the measured distance between the greater trochanters following Bennett et al. (82). The center of rotation of the ankle joint was assumed to lie at the anterior edge of the medial malleolus (83). During quiet standing, whole-body orientation was computed through a static inverted pendulum approximation, where the CoM was assumed to be directly above the CoP. The length of the pendulum was approximated as the center of mass height with respect to the ankle joint (*h*_CoM_), estimated by multiplying the body height by 0.56 for female and 0.57 for male participants (84), and subtracting the ankle height. The lean angle *θ* was then determined in radians by dividing the distance of the CoP with respect to the ankle joint (83) by the center of mass height.

Experiment 1 consisted of two trial types: preferred posture trials and target trials (Figure 1 C-E). In preferred posture trials, participants stood quietly facing forward with their arms at their sides, maintaining a comfortable posture with either their eyes open or closed. Each trial lasted five minutes. In target trials, participants were instructed to match their center of pressure (CoP) to a visual target displayed on a screen 3.20 meters in front of them, at a height of 1.67 meters. A black circle represented the target, and participants controlled a red cursor corresponding to their CoP location. The cursor turned green when it was within the target area, providing real-time feedback. While the target circle was always drawn in the center of the screen, participants had to maintain a lean angle *θ*_target_ of −2, 0, 2, 4, 6, or 10 · 10^-2^ rad to get the cursor in the center. These lean angles correspond to a subset of the targets used by Mensink, Nasrabadi (63), where the variability of ankle torques was investigated.

The corresponding CoP target positions were calculated as:

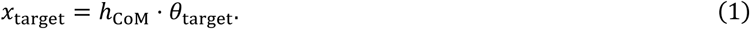

The radius *r*_target_ of the target circle was normalized to body length so that a similar whole-body angle excursion would result in a similar deviation from the target for all participants. The radius of the target was computed as

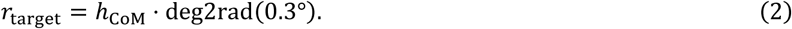

The circle was green when the participant was within the radius of the target, computed as

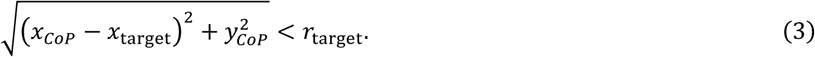

A 5-minute break was scheduled midway through the experiment. To minimize order effects, the order of the blocks (Preferred or Target) was reversed before and after the break. A total of 3 trials were excluded due to technical issues or experimenter error (EO and targets 0 and 2, all from different participants). In these cases, participant mean data were computed using the remaining valid trial.

#### Experiment 2: Walking initiation protocol

The aim of Experiment 2 was to determine the energetic cost of gait initiation across natural postures and to evaluate whether humans select postures that minimize this cost based on the expectation of walking direction. Participants (9 male, 11 female, Age = 24.1 ± 3.3 years, Height = 178.1 ± 11.1 cm, Weight = 70.0 ± 10.5 kg) stood on a force plate and initiated gait when a prompt on a screen asked them to start walking in the forwards or backwards direction (Figure 1F-G). Because the initiation of walking takes far less time than the time constant of indirect calorimetry (∼42 seconds (41)), the metabolic cost of gait initiation could not be measured. Instead, we estimated metabolic cost using musculoskeletal simulations on the kinetic and kinematic data measured through the force plate and motion capture system (see *Computation of metabolic cost* below).

The motion capture system consisted of 8 cameras running at 100 Hz (4 Miqus M3 and 4 Miqus M5, Qualisys AB, Göteborg, Sweden), an analog measurement board (Qualisys Analog Interface 16 Channels, 230597, Qualisys AB, Göteborg, Sweden), a sync box (Qualisys Sync Box, 410850, Qualisys AB, Göteborg, Sweden), and a single force plate running at 1000 Hz (Plate: BMS 400600HF-1K, Amplifier: Optima OPT-SC, AMTI, MA, US) embedded in a wooden walkway (740 cm long, 80 cm wide, >2mm clearance gap with the force plate and wooden walkway). The foot placement and motion capture marker locations were identical to those used in Experiment 1, with the addition of a marker on the T5 vertebra, which could now be placed since the back-worn metabolic analyzer was no longer used (see S1 Appendix for list of markers). A TV screen was placed at eye level at the anterior-facing end of the walkway (3.2 m away) to show the CoP target in some trials and to prompt the participant to start walking at the start of the trial. The data was stored in 12-second trials, with the prompt appearing at the 6-second mark.

Experiment 2 consisted of two sets of trials: preferred and target posture trials. The preferred posture set was always performed first to avoid influencing the participants’ preferred posture at gait initiation. In the preferred trials, the participant was instructed to stand calmly with their arms resting at their sides while looking straight ahead. During these trials, there were three distinct blocks to evaluate whether the known probability of gait initiation influenced posture: Forward-only (20 trials), Random (15 trials forward, 15 trials backward), and Backward-only (20 trials). In all three blocks, the participant was explicitly informed of the probabilities associated with the prompts, but not as to when they would be delivered. In the Random block, a random permutation determined the order of the prompts, and neither the participant nor the experimenter (to prevent experimenter bias) was aware of the direction presented for each trial.

In the set with target trials, participants received the same visual feedback on CoP position as used in Experiment 1 (Figure 1D, Eq 1-3), but only at target locations of −2, 2, and 6 · 10^-2^ rad. For each target, participants were prompted to walk forward or backward 5 times each, resulting in a total of 30 trials. The trials were ordered using a random permutation, ensuring that the target order and initiation direction were unpredictable. Trials that contained an error, such as initiating gait in the wrong direction or with the incorrect leg, were repeated and shuffled among the last five trials so that participants could not predict which direction would be repeated.

Prompts to initiate walking were provided using the TV screen. Forward trials were indicated by a blue screen displaying “Forward”, an upward-facing arrow, and a sound tone of 532 Hz; backward trials were indicated with an orange screen displaying “Backward”, a down-facing arrow, and a 440 Hz sound tone. After 3.1 seconds, a stop tone of 329 Hz was played to signal to participants that the trial had been completed. The period was selected to approximate the time by which participants should have made their fourth foot strike. Participants were asked to step with the same leg consistently and could choose a different leg for forward or backward walking. The time between a participant being ready and the delivery of a prompt was standardized to be a random integer in the range of 5-10 seconds, so that the timing of the prompt would not be predictable. The participant was considered ready when the time derivative of the center of pressure was below a threshold for 1000 samples (1000 ms) in a row. This threshold *ċ*_threshold_ was determined at the start of the experiment using a baseline trial of 2 minutes and computed as:

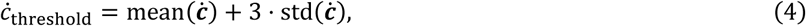

where *ċ* is the vector containing the center of pressure time derivative of the entire measurement. A MATLAB script was used to detect when the participant was ready and start a timer, indicating to the experimenter that the prompt could be sent.

### Quantification and statistical analysis

#### Data analysis

The average lean angle was extracted from center of pressure data using Equation 1. Further kinematic data were derived from musculoskeletal simulations (see below).

Raw EMG recordings were high-pass filtered (4th order zero-phase Butterworth, cutoff frequency 10 Hz), full-wave rectified, and then low-pass filtered (4th order zero-phase Butterworth, cutoff frequency 100 Hz). The erector spinae signal was high-pass filtered at 60 Hz instead to minimize electrocardiographic interference (85). The EMG signal for each muscle was normalized by dividing the signal by the maximum observed EMG signal during that participant’s maximum voluntary contraction exercises, which were additionally low-pass filtered at 6 Hz (79). The exercises consisted of prone knee flexion, supine ankle dorsiflexion, hip flexion kicks, maximum height jumps, and prone lumbar extension (79, 80).

Indirect calorimetry data were only used beyond the first 60 seconds of each recording, given that the time constant for this type of measurement is approximately 40 seconds (41). All metabolic measurements were normalized to body mass and reported in W/kg. The basal metabolic cost (i.e., the energy expended by non-motor processes to uphold bodily function) is generally assumed to be constant (86) and requires a prolonged period of rest and longer fasting to be measured accurately (32). Its contribution is commonly negated by subtracting the cost of a standing baseline trial from other activities, although this may underestimate the cost of those activities, as standing balance requires energy (86). In Experiment 1, the metabolic cost in the eyes-open preferred posture trial was subtracted from the other trials for both the measured and simulated energy expenditure separately for visualization purposes only. Doing so eliminated the offset between measured and simulated energy expenditure caused by the basal rate and muscle activity in the baseline trial, allowing for comparison of relative cost differences across different postures. Breath-by-breath samples lasting shorter than one second or having a respiratory quotient larger than 2 or smaller than 0.4 were omitted from the mean energy expenditure (0.69 % of samples). The 95% confidence interval of the mean energy expenditure was computed using bootstrapping by resampling the participant means 10,000 times and taking the 2.5th and 97.5th percentiles of the resulting distribution. Time-dependent measures of breathing (e.g., energy expenditure per second) were averaged using the breath duration for a weighted average, as a sample-based average would overvalue the contribution of short breaths.

In Experiment 2, the steady-state velocity of walking following gait initiation was determined as the maximum velocity of the center of mass between the presentation of the prompt and the end of the trial. The center of mass location was approximated using the inverse kinematics of the musculoskeletal model (see below). The time to reach steady state was determined as the time required to reach 90% of that participant’s average steady-state velocity. The time was reported in relation to the onset of movement to account for possible variations in reaction time. The onset of movement was determined as the first point in time where the magnitude of the center of pressure velocity in the lateral direction (due to the shifting of weight to one foot) exceeded the mean plus 3 times the standard deviation of the three seconds preceding the presentation of the prompt.

#### Musculoskeletal analysis of motion capture data

Motion capture trajectories were processed using Qualisys Track Manager (version 2024.2, Qualisys AB, Göteborg, Sweden). A prediction error of 20 mm and a maximal residual of 5 mm were accepted in the 3D tracking phase. Gaps in motion capture markers that can occur due to temporary occlusion were gap-filled using a polynomial interpolation if the occlusion lasted for at most 10 samples (100 ms). Markers that were occluded for longer than 100 ms were not gap-filled; in that segment of the measurement, only other valid markers were used for the subsequent inverse kinematics step (see below).

Musculoskeletal analyses were performed using OpenSim (13) (version 4.3) with the Rajagopal lower limb musculoskeletal model (12). First, the model was scaled to the participant’s anthropometrics using the OpenSim scale function, and an inverse kinematics analysis was performed to solve for the joint angles of the participants during standing and walking. Marker trajectories were low-pass filtered at 6 Hz. The weights used for the markers in the musculoskeletal scaling and inverse kinematics steps were empirically set, using the default settings (13, 87) as a starting point (values provided in S1 Appendix). To ensure that the foot was level with the ground for all participants after scaling, an ankle angle was prescribed per participant in the scaling step (5.24 ± 2.13°, weight 10) such that the foot of the resulting scaled model had a pitch angle close to 0° during standing. The headband markers were not used for inverse kinematics, as the head and torso were assumed to be a single rigid segment. For some female participants (Exp 1: 3/5, Exp 2: 5/11), the placement of the xiphoid process marker was hindered by anatomy and clothing, making it invisible to the motion capture cameras. In these cases, the marker was placed midway between the navel and the xiphoid process. This marker was assigned a low weight in all participants during scaling to accommodate the varied placement.

Inverse dynamics was then used to estimate the ground reaction forces. This estimation can be subtracted from the measured ground reaction forces to calculate the residual forces, serving as a metric for the simulation’s quality. Hicks et al. (43) recommend that the residual forces are less than 5% of the maximally measured net external force during that trial, and the residual moments less than 1% of the center of mass height times the maximally measured net external force. In Experiment 1, the median normalized residual, defined as the residual force or moment divided by the thresholds described above, was 0.101 (no unit) for residual forces and 0.363 for residual moments. The 95^th^ percentile of the normalized residual was 0.183 and 0.699, respectively, indicating that the simulations satisfy the recommended residual accuracy (43). In Experiment 2, the median force and moment badness were 0.182 and 0.431, with the 95^th^ percentile at 0.564 and 1.709. Here, the ground reaction forces were estimated using one force plate, which likely contributed to the normalized residual moments exceeding the threshold in some samples. Validation of this method is presented in the S1 Appendix.

Static muscle optimization was then performed at 20 Hz to estimate muscle activations by minimizing the squared activation (88–92). This approach estimates a set of muscle activations that can generate the required joint torques at each time point. Minimizing activation squared shares the load between muscles, but also assumes the ability to independently control single muscles without any neural constraints (for review, see Ting et al. (93)). The default implementation of the static optimization algorithm in OpenSim assumes (i) that the tendon is infinitely stiff, and (ii) that the parallel element of the Hill-type muscle does not contribute to the tensile force. However, passive stiffness around the ankle joint is known to be important during standing (94, 95). Therefore, the static optimization implementation by Uhlrich, Jackson (79), which includes the stiffnesses of the parallel element and the tendon, was used instead. This calculation utilizes the tendon stiffness provided in the musculoskeletal model (12) and optimizes the cost function using MATLAB.

To verify simulated muscle activity, it was compared to the EMG data of 7 major muscles contributing to postural control in Experiment 1. Hicks et al. (43) caution that EMG data should primarily serve to estimate the onset time of muscle activity and/or the trends between conditions, as data is hard to normalize and sensitive to measurement errors. Here, we compared the normalized EMG to the muscle activations to assess whether the increase in simulated muscle activity over lean angles matched the measured activation.

#### Computation of metabolic cost

The metabolic cost was estimated using the computed muscle activation from the OpenSim simulations. We used the metabolic energy model by Umberger et al. (30), which accounts for losses due to activation and maintenance, shortening, and mechanical work for each muscle independently. The activation and maintenance expenditure depend on the current fiber length of the muscle, with passive contributions reducing energy expenditure when the fiber length exceeds the optimal length (30). Fiber lengths were extracted from the OpenSim simulations, and optimal fiber lengths were derived from the model (12). Because fast-twitch (type II) fibers are metabolically less efficient than slow-twitch (type I) fibers (30, 52), each muscle’s energy cost was weighted by its ratio of slow-twitch muscle fibers. The muscle-specific fiber-type distributions were based on simulation code by Uchida et al. (96), who used data by Johnson, Polgar (29). Muscle mass was set using the values reported by Klein-Horsman et al. (31) and scaled proportionally to the body weight of each participant.

The Umberger model predicts a decrease in energy expenditure when muscle activation exceeds excitation. Unlike activation, excitation is not constrained by activation dynamics and is not computed in static muscle optimization methods as they assume idealized muscle activations without activation dynamics (89, 97, 98). Prior work shows that static methods produce similar results to dynamic methods (i.e., methods that compute excitation signals iteratively (99)) even during walking and running (89). Given the high computational cost of dynamic simulations (100), static optimization was deemed sufficient for modeling standing and gait initiation. The assumption that activation and excitation did not affect simulation results was verified by computing the maximum possible error between activation and excitation (See S1 Appendix).

The shortening heat rate was determined by identifying concentric versus eccentric contraction phases based on fiber length changes. Mechanical work was calculated as the product of muscle force and shortening velocity. The MATLAB analysis code, computation of energy expenditure, scaled OpenSim models, raw and analyzed data, and setup files for all OpenSim analyses performed were made publicly available (see Data availability statement).

#### Estimation of ground reaction forces from single force plate data

In Experiment 2, only one force plate was available for our experiments. To estimate the reaction forces under each foot, we combined kinematic data from foot markers with the measured global center of pressure (CoP). The CoP for each foot was assumed to lie along a line connecting the calcaneus (CAL) and second metatarsal (MT2) markers, projected onto the ground plane. The individual foot CoPs were estimated by identifying the shortest segment connecting this line to the measured global CoP, using constrained optimization. The normal force was proportionally distributed between the feet such that the moment in the ground plane was zero. This method was validated using an independent dataset with dual force plate measurements during gait initiation (see S1 Appendix for validation and detailed methods). Because reliable force estimates were only available before the first foot strike, static muscle optimization was performed only during this initial phase of gait initiation.

#### Monte-Carlo analysis of ankle stiffness and equilibrium lean angle

To investigate whether individual differences in passive ankle stiffness could account for variations in preferred lean angle (Experiment 1), we estimated how this variability affects the equilibrium whole-body lean angle. This analysis allows us to infer whether individual differences in the mechanical properties of the ankle could lead to different energetically optimal postural orientations. We performed a Monte Carlo simulation based on the parameter distributions for ankle stiffness reported by Moseley, Crosbie (42). The equilibrium lean angle was defined as the whole-body angle at which the moment produced by passive ankle stiffness balanced the gravitational moment acting on the body. The mass-normalized ankle stiffness of one ankle was modeled as

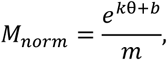

Where *θ* is lean angle, *k* = 56.09 ± 8.36, *b* = 1.871 ± 0.282, and *m* = 68.5 kg. These values were obtained from the study (42) and normalized by dividing the ankle moment by the average mass. We generated 50,000 samples from the distributions of *k* and *b*, assuming a normal distribution for both. Next, we computed the equilibrium angle *θ* for each combination by finding the intersection of the mass-normalized ankle moment (*M*_norm_(*θ*)) and gravitational moment (*gLθ*) using MATLAB for an assumed body with the mean parameters of the participants in Experiment 1 (Center of mass height = 0.954 m, weight not required). Because the resulting distribution of equilibrium angles was skewed, we report the median and interquartile range of the estimated equilibrium angles. The code implementation of the Monte Carlo analysis was made publicly available (see Data availability statement).

#### Statistical tests

Statistical analyses were performed in JASP (101) (rmANOVA) and MATLAB (*t*-tests), using a significance level of 0.05. Variables are expressed using mean ± standard deviation unless mentioned otherwise. Data for the Monte Carlo analysis were reported as median and interquartile range to account for the non-normality of the distribution. Degrees of freedom are reported between brackets for t-tests and rmANOVA outcomes. When sphericity was violated, Greenhouse-Geisser corrections were applied.

In Experiment 1, we used one-way repeated-measures ANOVAs across the six target conditions to determine whether whole-body orientation influenced various metrics of posture, including energy expenditure (measured and simulated), postural variability (center of pressure velocity and standard deviation of lean angle), and breathing characteristics (minute volume and breathing rate). If the main effect was significant, post-hoc comparisons between targets were conducted using paired-samples *t*-tests with Holm correction. For energy expenditure, post-hoc tests were performed by assessing which targets imposed a larger cost than the minimum-cost posture (2 · 10^-2^ rad, see Results) using one-sided paired-samples *t*-tests with Holm correction. To assess the energetic cost increase in the anterior and posterior directions, we used a paired-samples *t*-test to compare the average slopes. Additionally, we compared the energetic costs in the preferred posture trials with eyes open and eyes closed using a paired-samples *t*-test to assess the role of vision. Lastly, the eyes-open preferred trial was compared to all six targets using paired-samples *t*-tests with Holm correction.

In Experiment 2, we analyzed the lean angle prior to gait initiation along with various additional gait initiation metrics (e.g., energy expended and maximum velocity). To evaluate whether participants adjusted their posture or gait initiation characteristics when the walking direction was known, we compared the Random block to the Forward-only and Backward-only blocks using paired-samples *t*-tests. Additionally, we compared the Forward-only and Backward-only blocks using paired-samples *t*-tests to assess differences in body orientation and gait characteristics of forward and backward gait. To examine the effect of the three prescribed lean angles on gait initiation, we analyzed the target posture trials using separate one-way repeated measures ANOVAs for forward and backward gait initiations. Forward and backward initiations were analyzed separately because the interpretation of a given target depends on the walking direction. For example, a backwards-leaning posture is congruent for backwards but incongruent for forward initiation, meaning the interpretation of the lean angle is dependent on the lean angle. This violates the assumption of independence of repeated measures factors required for a two-way ANOVA. Post-hoc comparisons between lean angles were performed using Holm-corrected paired-samples t-tests.

## Acknowledgments

The authors would like to thank Timo Vanderhave and Paul de Jongh for their contributions to the preparation and data collection of Experiment 1, as well as Koen Jongbloed, Natasha Giri, and Ragnhild Maarleveld for their aid in preparing equipment for Experiment 1. We thank Daphne Onderwater for her assistance in labeling the markers for Experiment 2. We thank Matthew Millard for bringing the current force plate decomposition method to our attention. We thank Eline van der Kruk and Katla Guðmundsdóttir for providing access to the unpublished stand-to-walk trial data used to validate the ground reaction force decomposition method. Lastly, we would like to thank Johan Pel, Maarten Frens, and Lucas Mensink for their help in the ideation and conceptualization phase of this study.

## Author contributions

Conceptualization: M.L., A.S., P.A.F.

Data Curation: M.L., P.A.F.

Formal Analysis: M.L.

Funding Acquisition: P.A.F.

Investigation: M.L., N.v.A., P.A.F.

Methodology: M.L., N.v.A., A.S., P.A.F.

Project Administration: P.A.F.

Resources: A.S., P.A.F.

Software: M.L.

Supervision: M.L., A.S., P.A.F.

Validation: M.L., A.S., P.A.F.

Visualization: M.L.

Writing – Original Draft: M.L., P.A.F.

Writing – Review & Editing: M.L., N.v.A., A.S., P.A.F.

## Supplementary material

**S1 Table. Experiment 1 Group Data.** Data table containing participant averages and standard deviations of various metrics in the target and preferred posture conditions of Experiment 1.

**S2 Table. Experiment 2 Group Data.** Data table containing participant averages and standard deviations of various metrics in the gait initiation trials using the target or preferred posture conditions of Experiment 2.

**S1 Appendix. Supplemental Methods and Validation of Assumptions.** Contains validation of the force plate splitting algorithm, validation of the indirect calorimetry trial length, validation of the assumption that excitation and excitation are similar in slow movements, and a list of used motion capture markers and their weights for scaling and inverse kinematics simulations.

## Data availability statement

All data and code supporting the findings of this study are publicly available in “Data and code for *‘The energetic cost of human standing balance and gait initiation over a range of natural postures’*”, [doi link], DataverseNL. Specifically, the shared data included preprocessed motion capture (.mat and .trc), ground reaction force (.mat and .sto), electromyography (Experiment 1 only, .mat), and metabolic analyzer data (Experiment 1 only, .csv and .mat). OpenSim simulations and setup files (.xml) are provided for the scaling (.osim model per participant), inverse kinematics (.mot), inverse dynamics (.sto), and static optimization (performed in MATLAB, .sto or .csv). Provided code includes: MATLAB code for pre-processing data (.mat), executing all listed OpenSim analyses, analysis necessary to generate the presented figures, a custom MATLAB interface with Qualisys Track Manager for presenting the center of pressure target, code for EMG acquisition based on the TMSi MATLAB interface, and the Monte Carlo simulation for passive ankle moment.

## References

1. Selinger JC, O’Connor SM, Wong JD, Donelan JM. Humans Can Continuously Optimize Energetic Cost during Walking. Curr Biol. 2015;25(18):2452–6.

2. Selinger JC, Wong JD, Simha SN, Donelan JM. How humans initiate energy optimization and converge on their optimal gaits. J Exp Biol. 2019;222(Pt 19).

3. Abram SJ, Poggensee KL, Sanchez N, Simha SN, Finley JM, Collins SH, et al. General variability leads to specific adaptation toward optimal movement policies. Curr Biol. 2022;32(10):2222–32 e5.

4. Carlisle RE, Kuo AD. Optimization of energy and time predicts dynamic speeds for human walking. Elife. 2023;12.

5. Felt W, Selinger JC, Donelan JM, Remy CD. “Body-In-The-Loop”: Optimizing Device Parameters Using Measures of Instantaneous Energetic Cost. PLOS ONE. 2015;10(8):e0135342.

6. Rathkey JK, Wall-Scheffler CM. People choose to run at their optimal speed. Am J Phys Anthropol. 2017;163(1):85–93.

7. Bertram JE. Locomotion: Why We Walk the Way We Walk. Curr Biol. 2015;25(18):R795–7.

8. Miller RH, Umberger BR, Hamill J, Caldwell GE. Evaluation of the minimum energy hypothesis and other potential optimality criteria for human running. Proceedings of the Royal Society B: Biological Sciences. 2012;279(1733):1498–505.

9. Zelik KE, Kuo AD. Human walking isn’t all hard work: evidence of soft tissue contributions to energy dissipation and return. Journal of Experimental Biology. 2010;213(24):4257–64.

10. Tucker VA. Energetic cost of locomotion in animals. Comp Biochem Physiol. 1970;34(4):841–6.

11. Kuo AD, Donelan JM, Ruina A. Energetic consequences of walking like an inverted pendulum: step-to-step transitions. Exerc Sport Sci Rev. 2005;33(2):88–97.

12. Rajagopal A, Dembia CL, DeMers MS, Delp DD, Hicks JL, Delp SL. Full-Body Musculoskeletal Model for Muscle-Driven Simulation of Human Gait. IEEE Trans Biomed Eng. 2016;63(10):2068–79.

13. Seth A, Hicks JL, Uchida TK, Habib A, Dembia CL, Dunne JJ, et al. OpenSim: Simulating musculoskeletal dynamics and neuromuscular control to study human and animal movement. PLoS Comput Biol. 2018;14(7):e1006223.

14. Uchida TK, Seth A, Pouya S, Dembia CL, Hicks JL, Delp SL. Simulating Ideal Assistive Devices to Reduce the Metabolic Cost of Running. PLoS One. 2016;11(9):e0163417.

15. Poggensee KL, Collins SH. Lower limb biomechanics of fully trained exoskeleton users reveal complex mechanisms behind the reductions in energy cost with human-in-the-loop optimization. Frontiers in Robotics and AI. 2024;11.

16. Uchida TK, Delp SL, Delp D. Biomechanics of Movement: The Science of Sports, Robotics, and Rehabilitation: MIT Press; 2021.

17. Smith JW. The forces operating at the human ankle joint during standing. J Anat. 1957;91(4):545–64.

18. Nashner LM. Adapting reflexes controlling the human posture. Exp Brain Res. 1976;26(1):59–72.

19. Gage WH, Winter DA, Frank JS, Adkin AL. Kinematic and kinetic validity of the inverted pendulum model in quiet standing. Gait Posture. 2004;19(2):124–32.

20. Fitzpatrick RC, Taylor JL, McCloskey DI. Ankle stiffness of standing humans in response to imperceptible perturbation: reflex and task-dependent components. J Physiol. 1992;454:533–47.

21. Kuo AD. An optimal control model for analyzing human postural balance. IEEE Trans Biomed Eng. 1995;42(1):87–101.

22. Lockhart DB, Ting LH. Optimal sensorimotor transformations for balance. Nat Neurosci. 2007;10(10):1329–36.

23. Ito S, Saka Y, Kawasaki H. Where center of pressure should be controlled in biped upright posture. Systems and Computers in Japan. 2004;35(5):23–31.

24. van der Kooij H, Jacobs R, Koopman B, Grootenboer H. A multisensory integration model of human stance control. Biol Cybern. 1999;80(5):299–308.

25. Shanbhag J, Fleischmann S, Wechsler I, Gassner H, Winkler J, Eskofier BM, et al. A sensorimotor enhanced neuromusculoskeletal model for simulating postural control of upright standing. Frontiers in Neuroscience. 2024;18.

26. Suzuki Y, Geyer H. A Neuro-Musculo-Skeletal Model of Human Standing Combining Muscle-Reflex Control and Virtual Model Control. Annu Int Conf IEEE Eng Med Biol Soc. 2018;2018:5590–3.

27. Forbes PA, Chen A, Blouin JS. Sensorimotor control of standing balance. Handb Clin Neurol. 2018;159:61–83.

28. Kiemel T, Zhang Y, Jeka JJ. Identification of neural feedback for upright stance in humans: stabilization rather than sway minimization. J Neurosci. 2011;31(42):15144–53.

29. Johnson MA, Polgar J, Weightman D, Appleton D. Data on the distribution of fibre types in thirty-six human muscles: An autopsy study. Journal of the Neurological Sciences. 1973;18(1):111–29.

30. Umberger BR, Gerritsen KGM, Martin PE. A Model of Human Muscle Energy Expenditure. Computer Methods in Biomechanics and Biomedical Engineering. 2003;6(2):99–111.

31. Klein Horsman MD, Koopman HF, van der Helm FC, Prose LP, Veeger HE. Morphological muscle and joint parameters for musculoskeletal modelling of the lower extremity. Clin Biomech (Bristol, Avon). 2007;22(2):239–47.

32. Amaro-Gahete FJ, Sanchez-Delgado G, Alcantara JMA, Martinez-Tellez B, Acosta FM, Merchan-Ramirez E, et al. Energy expenditure differences across lying, sitting, and standing positions in young healthy adults. PLOS ONE. 2019;14(6):e0217029.

33. Judice PB, Hamilton MT, Sardinha LB, Zderic TW, Silva AM. What is the metabolic and energy cost of sitting, standing and sit/stand transitions? Eur J Appl Physiol. 2016;116(2):263–73.

34. Betts JA, Smith HA, Johnson-Bonson DA, Ellis TI, Dagnall J, Hengist A, et al. The Energy Cost of Sitting versus Standing Naturally in Man. Medicine & Science in Sports & Exercise. 2019;51(4):726–33.

35. Saeidifard F, Medina-Inojosa JR, Supervia M, Olson TP, Somers VK, Erwin PJ, et al. Differences of energy expenditure while sitting versus standing: A systematic review and meta-analysis. Eur J Prev Cardiol. 2018;25(5):522–38.

36. Houdijk H, Fickert R, van Velzen J, van Bennekom C. The energy cost for balance control during upright standing. Gait Posture. 2009;30(2):150–4.

37. Houdijk H, Brown SE, van Dieen JH. Relation between postural sway magnitude and metabolic energy cost during upright standing on a compliant surface. J Appl Physiol (1985). 2015;119(6):696–703.

38. Saha D, Gard S, Fatone S, Ondra S. The effect of trunk-flexed postures on balance and metabolic energy expenditure during standing. Spine (Phila Pa 1976). 2007;32(15):1605–11.

39. Orendurff MS. How humans walk: Bout duration, steps per bout, and rest duration. The Journal of Rehabilitation Research and Development. 2008;45(7):1077–90.

40. Fawver B, Roper JA, Sarmento C, Hass CJ. Forward leaning alters gait initiation only at extreme anterior postural positions. Human Movement Science. 2018;59:1–11.

41. Selinger JC, Donelan JM. Estimating instantaneous energetic cost during non-steady-state gait. J Appl Physiol (1985). 2014;117(11):1406–15.

42. Moseley AM, Crosbie J, Adams R. Normative data for passive ankle plantarflexion--dorsiflexion flexibility. Clin Biomech (Bristol, Avon). 2001;16(6):514–21.

43. Hicks JL, Uchida TK, Seth A, Rajagopal A, Delp SL. Is my model good enough? Best practices for verification and validation of musculoskeletal models and simulations of movement. J Biomech Eng. 2015;137(2):020905.

44. Le Mouel C, Brette R. Mobility as the Purpose of Postural Control. Front Comput Neurosci. 2017;11:67.

45. Zhao G, Grimmer M, Seyfarth A. The mechanisms and mechanical energy of human gait initiation from the lower-limb joint level perspective. Scientific Reports. 2021;11(1).

46. Miller CA, Verstraete MC. A mechanical energy analysis of gait initiation. Gait Posture. 1999;9(3):158–66.

47. Camargo J, Ramanathan A, Flanagan W, Young A. A comprehensive, open-source dataset of lower limb biomechanics in multiple conditions of stairs, ramps, and level-ground ambulation and transitions. Journal of Biomechanics. 2021;119:110320.

48. Anand M, Seipel J, Rietdyk S. A modelling approach to the dynamics of gait initiation. Journal of The Royal Society Interface. 2017;14(128):20170043.

49. Wade MG, Jones G. The role of vision and spatial orientation in the maintenance of posture. Phys Ther. 1997;77(6):619–28.

50. Rasman BG, Forbes PA, Tisserand R, Blouin JS. Sensorimotor Manipulations of the Balance Control Loop-Beyond Imposed External Perturbations. Front Neurol. 2018;9:899.

51. Rodman PS, McHenry HM. Bioenergetics and the origin of hominid bipedalism. Am J Phys Anthropol. 1980;52(1):103–6.

52. O’Neill MC, Umberger BR, Holowka NB, Larson SG, Reiser PJ. Chimpanzee super strength and human skeletal muscle evolution. Proceedings of the National Academy of Sciences. 2017;114(28):7343–8.

53. Raichlen DA, Armstrong H, Lieberman DE. Calcaneus length determines running economy: implications for endurance running performance in modern humans and Neandertals. J Hum Evol. 2011;60(3):299–308.

54. Sinha T, Maki BE. Effect of forward lean on postural ankle dynamics. IEEE Trans Rehabil Eng. 1996;4(4):348–59.

55. Gottlieb GL, Agarwal GC. Response to sudden torques about ankle in man: myotatic reflex. J Neurophysiol. 1979;42(1 Pt 1):91–106.

56. Hof AL, Gazendam MG, Sinke WE. The condition for dynamic stability. J Biomech. 2005;38(1):1–8.

57. Horak FB. Postural orientation and equilibrium: what do we need to know about neural control of balance to prevent falls? Age Ageing. 2006;35 Suppl 2:ii7–ii11.

58. Pozzo T, Berthoz A, Lefort L. Head stabilization during various locomotor tasks in humans. Experimental Brain Research. 1990;82(1).

59. Wickens CD. Multiple resources and mental workload. Hum Factors. 2008;50(3):449–55.

60. Boisgontier MP, Beets IAM, Duysens J, Nieuwboer A, Krampe RT, Swinnen SP. Age-related differences in attentional cost associated with postural dual tasks: Increased recruitment of generic cognitive resources in older adults. Neuroscience & Biobehavioral Reviews. 2013;37(8):1824–37.

61. Carpenter MG, Murnaghan CD, Inglis JT. Shifting the balance: evidence of an exploratory role for postural sway. Neuroscience. 2010;171(1):196–204.

62. Woodhull AM, Maltrud K, Mello BL. Alignment of the human body in standing. Eur J Appl Physiol Occup Physiol. 1985;54(1):109–15.

63. Mensink LH, Nasrabadi AM, Rasman BG, Blouin J-S, Forbes PA. The sense and control of standing balance in the presence of motor noise. (in preparation). 2025.

64. Duarte M, Zatsiorsky VM. On the fractal properties of natural human standing. Neurosci Lett. 2000;283(3):173–6.

65. Duarte M, Zatsiorsky VM. Patterns of center of presure migration during prolonged unconstrained standing. Motor Control. 1999;3(1):12–27.

66. Asai Y, Tasaka Y, Nomura K, Nomura T, Casadio M, Morasso P. A model of postural control in quiet standing: robust compensation of delay-induced instability using intermittent activation of feedback control. PLoS One. 2009;4(7):e6169.

67. Loram ID, van de Kamp C, Lakie M, Gollee H, Gawthrop PJ. Does the motor system need intermittent control? Exerc Sport Sci Rev. 2014;42(3):117–25.

68. Suzuki Y, Nomura T, Casadio M, Morasso P. Intermittent control with ankle, hip, and mixed strategies during quiet standing: a theoretical proposal based on a double inverted pendulum model. J Theor Biol. 2012;310:55–79.

69. Huryn TP, Blouin J-S, Croft EA, Koehle MS, Van Der Loos HFM. Experimental Performance Evaluation of Human Balance Control Models. IEEE Transactions on Neural Systems and Rehabilitation Engineering. 2014;22(6):1115–27.

70. Kaminishi K, Jiang P, Chiba R, Takakusaki K, Ota J. Postural control of a musculoskeletal model against multidirectional support surface translations. PLOS ONE. 2019;14(3):e0212613.

71. Jiang P, Chiba R, Takakusaki K, Ota J. Generation of the Human Biped Stance by a Neural Controller Able to Compensate Neurological Time Delay. PLOS ONE. 2016;11(9):e0163212.

72. Afschrift M, Jonkers I, De Schutter J, De Groote F. Mechanical effort predicts the selection of ankle over hip strategies in nonstepping postural responses. J Neurophysiol. 2016;116(4):1937–45.

73. Ding Z, Tsang CK, Nolte D, Kedgley AE, Bull AMJ. Improving Musculoskeletal Model Scaling Using an Anatomical Atlas: The Importance of Gender and Anthropometric Similarity to Quantify Joint Reaction Forces. IEEE Transactions on Biomedical Engineering. 2019;66(12):3444–56.

74. Maarleveld R, Veeger H, van der Helm F, Son J, Lieber R, van der Kruk E. What the% PCSA? Addressing Diversity in Lower-Limb Musculoskeletal Models: Age-and Sex-related Differences in PCSA and Muscle Mass. arXiv preprint arXiv:241100071. 2024.

75. Dos Santos DA, Fukuchi CA, Fukuchi RK, Duarte M. A data set with kinematic and ground reaction forces of human balance. PeerJ. 2017;5:e3626.

76. Brockway JM. Derivation of formulae used to calculate energy expenditure in man. Hum Nutr Clin Nutr. 1987;41(6):463–71.

77. Visser M, Deurenberg P, van Staveren WA, Hautvast JG. Resting metabolic rate and diet-induced thermogenesis in young and elderly subjects: relationship with body composition, fat distribution, and physical activity level. Am J Clin Nutr. 1995;61(4):772–8.

78. Hermens HJ, Freriks B, Disselhorst-Klug C, Rau G. Development of recommendations for SEMG sensors and sensor placement procedures. J Electromyogr Kinesiol. 2000;10(5):361–74.

79. Uhlrich SD, Jackson RW, Seth A, Kolesar JA, Delp SL. Muscle coordination retraining inspired by musculoskeletal simulations reduces knee contact force. Scientific Reports. 2022;12(1):9842.

80. Suydam SM, Manal K, Buchanan TS. The Advantages of Normalizing Electromyography to Ballistic Rather than Isometric or Isokinetic Tasks. J Appl Biomech. 2017;33(3):189–96.

81. Hagströmer M, Oja P, Sjöström M. The International Physical Activity Questionnaire (IPAQ): a study of concurrent and construct validity. Public Health Nutrition. 2006;9(6):755–62.

82. Bennett HJ, Shen G, Weinhandl JT, Zhang S. Validation of the greater trochanter method with radiographic measurements of frontal plane hip joint centers and knee mechanical axis angles and two other hip joint center methods. J Biomech. 2016;49(13):3047–51.

83. Sado N, Shiotani H, Saeki J, Kawakami Y. Positional difference of malleoli-midpoint from three-dimensional geometric centre of rotation of ankle and its effect on ankle joint kinetics. Gait Posture. 2021;83:223–9.

84. Virmavirta M, Isolehto J. Determining the location of the body׳s center of mass for different groups of physically active people. J Biomech. 2014;47(8):1909–13.

85. Redfern M, Hughes R, Chaffin D. High-pass filtering to remove electrocardiographic interference from torso EMG recordings. Clinical Biomechanics. 1993;8(1):44–8.

86. Weyand PG, Smith BR, Sandell RF, editors. Assessing the metabolic cost of walking: The influence of baseline subtractions. 2009 Annual International Conference of the IEEE Engineering in Medicine and Biology Society; 2009 3-6 Sept. 2009.

87. Delp SL, Loan JP, Hoy MG, Zajac FE, Topp EL, Rosen JM. An interactive graphics-based model of the lower extremity to study orthopaedic surgical procedures. IEEE Trans Biomed Eng. 1990;37(8):757–67.

88. Crowninshield RD, Brand RA. A physiologically based criterion of muscle force prediction in locomotion. J Biomech. 1981;14(11):793–801.

89. Anderson FC, Pandy MG. Static and dynamic optimization solutions for gait are practically equivalent. J Biomech. 2001;34(2):153–61.

90. Erdemir A, Mclean S, Herzog W, Van Den Bogert AJ. Model-based estimation of muscle forces exerted during movements. Clinical Biomechanics. 2007;22(2):131–54.

91. Herzog W, Binding P. Predictions of antagonistic muscular activity using nonlinear optimization. Math Biosci. 1992;111(2):217–29.

92. Prilutsky BI, Herzog W, Allinger TL. Forces of individual cat ankle extensor muscles during locomotion predicted using static optimization. Journal of Biomechanics. 1997;30(10):1025–33.

93. Ting LH, Chvatal SA, Safavynia SA, Lucas Mckay J. Review and perspective: neuromechanical considerations for predicting muscle activation patterns for movement. International Journal for Numerical Methods in Biomedical Engineering. 2012;28(10):1003–14.

94. Amankwah K, Triolo R, Kirsch R, Audu M. A model-based study of passive joint properties on muscle effort during static stance. J Biomech. 2006;39(12):2253–63.

95. Sakanaka TE, Gill J, Lakie MD, Reynolds RF. Intrinsic ankle stiffness during standing increases with ankle torque and passive stretch of the Achilles tendon. PLOS ONE. 2018;13(3):e0193850.

96. Uchida TK, Hicks JL, Dembia CL, Delp SL. Stretching Your Energetic Budget: How Tendon Compliance Affects the Metabolic Cost of Running. PLOS ONE. 2016;11(3):e0150378.

97. Thelen DG, Anderson FC, Delp SL. Generating dynamic simulations of movement using computed muscle control. J Biomech. 2003;36(3):321–8.

98. Dembia CL, Bianco NA, Falisse A, Hicks JL, Delp SL. OpenSim Moco: Musculoskeletal optimal control. PLoS Comput Biol. 2020;16(12):e1008493.

99. Thelen DG, Anderson FC. Using computed muscle control to generate forward dynamic simulations of human walking from experimental data. J Biomech. 2006;39(6):1107–15.

100. Belli I, Joshi S, Prendergast JM, Beck I, Della Santina C, Peternel L, et al. Does enforcing glenohumeral joint stability matter? A new rapid muscle redundancy solver highlights the importance of non-superficial shoulder muscles. PLOS ONE. 2023;18(11):e0295003.

101. JASP Team. JASP (Version 0.18.1) [Computer software]. 2023.

